# Set1-mediated histone H3K4 methylation is required for azole induction of the ergosterol biosynthesis genes and antifungal drug resistance in *Candida glabrata*

**DOI:** 10.1101/2021.11.17.469015

**Authors:** Kortany M. Baker, Smriti Hoda, Debasmita Saha, Livia Georgescu, Nina D. Serratore, Yueping Zhang, Nadia A. Lanman, Scott D. Briggs

**Author notes:** To whom correspondence should be addressed: Scott D. Briggs, Department of Biochemistry Hansen Life Science Research Building, 201 S. University Street, West Lafayette, IN 47907. Footnote: Current address for Yueping Zhang and Nina Serratore Nina Serratore: Cook Research Incorporated, 1 Geddes Way, West Lafayette, IN 47906 Yueping Zhang: College of Veterinary Medicine, China Agricultural University, No. 2 Yuanmingyuan West Road, Haidian District, Beijing 100193, China.

## Abstract

*Candida glabrata* is an opportunistic pathogen that has developed the ability to adapt and thrive under azole treated conditions. The common mechanisms that can result in *Candida* drug resistance are due to mutations or overexpression of the drug efflux pump or the target of azole drugs, Cdr1 and Erg11, respectively. However, the role of epigenetic histone modifications in azole-induced gene expression and drug resistance are poorly understood in *C. glabrata*. In this study, we show for the first time that Set1 mediates histone H3K4 mono-, di-, and trimethylation in *C. glabrata*. In addition, loss of *SET1* and histone H3K4 methylation results in increased susceptibility to azole drugs in both *C. glabrata* and *S. cerevisiae*. Intriguingly, this increase in susceptibility to azole drugs in strains lacking Set1-mediated histone H3K4 methylation is not due to altered transcript levels of *CDR1*, *PDR1* or Cdr1’s ability to efflux drugs. Genome-wide transcript analysis revealed that Set1 is necessary for azole-induced expression of 12 genes involved in the late biosynthesis of ergosterol including *ERG11* and *ERG3*. Importantly, chromatin immunoprecipitation analysis showed that histone H3K4 trimethylation was detected on chromatin of actively transcribed *ERG* genes. Furthermore, H3K4 trimethylation increased upon azole-induced gene expression which was also found to be dependent on the catalytic activity of Set1. Altogether, our findings show that Set1-mediated histone H3K4 methylation governs the intrinsic drug resistant status in *C. glabrata* via epigenetic control of azole-induced *ERG* gene expression.

**IMPORTANCE:** *C. glabrata* is the second most commonly isolated species from *Candida* infections, coming in second to *C. albicans*. Treatment of *C. glabrata* infections are difficult due to their natural resistance to antifungal azole drugs and their ability to adapt and become multidrug resistant. In this study, we investigated the contributing cellular factors for controlling drug resistance. We have determined that an epigenetic mechanism governs the expression of genes involved in the late ergosterol biosynthesis pathway, an essential pathway that antifungal drugs target. This epigenetic mechanism involves histone H3K4 methylation catalyzed by the Set1 methyltransferase complex (COMPASS). We also show that Set1-mediated histone H3K4 methylation is needed for expression of specific azole induced genes and azole drug resistance in *C. glabrata*. Identifying epigenetic mechanisms contributing to drug resistance and pathogenesis could provide alternative targets for treating patients with fungal infections.

## INTRODUCTION

*Candida* infections are a major health concern due to the increased frequency of infections and the development of drug resistance (1, 2). Over the years, *Candida glabrata* has become the second most common cause of candidiasis (1–3). In some immunocompromised patients, such as diabetics, patients with hematologic cancer, organ transplant recipients, and the elderly, it is the most predominate *Candida* infection (2–6). The emergence of *C. glabrata* as a major pathogen is likely due to its intrinsic drug resistance to azole antifungal drugs and ability to quickly adapt and acquire clinical drug resistance during treatment (3, 7). The consequence of drug resistance leads to increases in healthcare costs as well as lower success rates in treatment and an increase in mortality (8–10).

*C. glabrata* naturally has low susceptibility to azole drugs and because of this attribute, echinocandins are the preferred drug choice for treating *C. glabrata* infections (11). *C. glabrata* can also acquire clinical resistance to azole drugs which is often due to overexpressing the ABC-transporter drug efflux pump Cdr1 or Pdh1 (Cdr2) caused by gain of function mutations in the transcription factor Pdr1 (7, 12–14). In other *Candida* species, acquired clinical azole resistance can also be due to overexpression of *ERG11* due to gain of function mutations in the Upc2 transcription factor or mutations in *ERG11* (15–17). However, for unbeknownst reasons, *ERG11* or *UPC2* mutations are typically not found in clinically drug resistant *C. glabrata* strains (7, 18–20).

Because pathogenic fungi can rapidly adapt to various cellular environments and xenobiotic drug exposures, epigenetic mechanisms are also likely contributing to altered gene expression profiles permissive for adaptation and drug resistance. Several studies in *C. albicans* support this hypothesis and show that epigenetic factors such as histone acetyltransferases, *Ca*Gcn5 and *Ca*Rtt109, and histone deacetylases, *Ca*Rpd3 and *Ca*Hda1 are important for either fungal pathogenesis and/or drug resistance (21–25). In contrast, epigenetic factors that post-translationally modify histones have not been extensively studied for their roles in drug resistance in *C. glabrata*. Nonetheless, Orta- Zavalz et al., have shown that deleting histone deacetylase, *CgHST1* decreases susceptibility to fluconazole which is likely attributed to an increase in transcript levels of *CgPDR1* and *CgCDR1* under untreated conditions (26). In addition, a recent publication by Filler et al., has indicated that *C. glabrata* strains that are deleted for *GCN5*, *RPD3*, or *SPP1* have increased susceptibility to caspofungin when using high concentrations (27). However, no mechanistic understanding such as gene targets or changes in chromatin/histone modifications was provided for the caspofungin hypersensitive phenotype.

Previous publications from our lab demonstrated that in *S. cerevisiae* loss of Set1, a known histone H3K4 methyltransferase, has a hypersensitive growth defect in the presence of the antifungal metabolite, brefeldin A (BFA) and clinically used azole drugs (28, 29). We determined that hypersensitivity to BFA was due to a decrease in ergosterol levels in *S. cerevisiae* strains lacking histone H3K4 methylation. However, until this study, no mechanistic understanding has been provided why a strain lacking *SET1* alters azole drug susceptibility. Furthermore, in *C. albicans*, loss of *SET1* appears to alter virulence but not azole drug resistance (30). To determine if an increase in azole susceptibility is conserved in a human fungal pathogen closely related to *S. cerevisiae*, we investigated the role of Set1 and its mechanistic contribution to drug resistance in *C. glabrata*.

In this study, we show for the first time that Set1-mediates histone H3K4 mono-, di-, and trimethylation in *C. glabrata* and loss of Set1-mediated histone H3K4 methylation alters the azole drug susceptibility of *C. glabrata* similar to what is seen in *S. cerevisiae*. This increase in susceptibility to azole drugs in *C. glabrata* strains lacking Set1-mediated histone H3K4 methylation is not a consequence of altered expression levels of *CDR1*, *PDR1* or their ability to efflux drugs. Interestingly, RNA-sequencing (RNA-seq) revealed that Set1 is required for azole-induced expression of *ERG* genes, including *ERG11* and *ERG3.* This azole-induced gene expression was dependent on Set1 methyltransferase activity and associated with gene-specific increases in histone H3K4 trimethylation on *ERG11* and *ERG3* chromatin. Overall, we have provided a mechanistic understanding of why Set1 mediated histone H3K4 methylation governs the intrinsic drug resistant status in *C. glabrata.* Identifying and understanding the epigenetic mechanisms contributing to drug resistance will be important for the development of alternative drug targets for treating patients with fungal infections.

## RESULTS

### Loss of Set1-mediated histone H3K4 methylation in *S. cerevisiae* and *C. glabrata* alters azole drug efficacy

Set1 is a known SET domain-containing lysine histone methyltransferase that is conserved from yeast to humans and the enzymatic activity of the SET domain catalyzes mono-, di-, and trimethylation on histone H3 at Lysine 4 (Lys4) (31, 32). Our previous work in *Saccharomyces cerevisiae* (*S. cerevisiae*) has determined that loss of *SET1* in the BY4741 background strain results in increased susceptibility to azole drugs suggesting that H3K4 methylation is necessary for mediating wild-type azole drug resistance. To determine the role of histone H3K4 methylation in azole drug efficacy, we constructed histone H3K4R mutations in the BY4741 background strain. Because *S. cerevisiae* has two genes encoding histone H3, two yeast strains were constructed where a histone H3K4R mutation was integrated at one histone H3 gene keeping the other gene wild-type (ScH3K4R-1) while the other strain contained H3K4R mutations integrated at both histone H3 genes (ScH3K4R-2, see supplemental table S1). To determine if loss of histone H3K4 methylation altered azole drug sensitivity similar to a *set1Δ* (Sc*set1Δ*) strain, a serial-dilution spot assay was performed. Both Sc*set1Δ* and *Sc*H3K4R mutant strains were grown in synthetic complete minimal media and spotted on SC agar plates with and without 8 µg/mL fluconazole (Fig. 1A). These data show that loss of histone H3 methylation by deleting Sc*SET1* or mutating histone H3 where both histone H3 genes are mutated at K4 (ScH3K4R-2), resulted in similar azole drug hypersensitivity when compared to each other (Fig. 1A). To confirm that histone H3K4 methylation was abolished in these strains, western blot analysis was performed using methyl-specific antibodies to detect histone H3K4 mono-, di-, and trimethylation (Fig. 1B). Histone H3 was used for a loading control (Fig. 1B). As expected, histone methylation was abolished in *set1Δ* and in H3K4R-2 mutation strains but not in the histone H3K4R-1 strain (Fig. 1B). Together our data demonstrate that the presence of Histone H3K4 methylation is critical for maintaining wild-type azole drug susceptibility.

**FIG 1.**
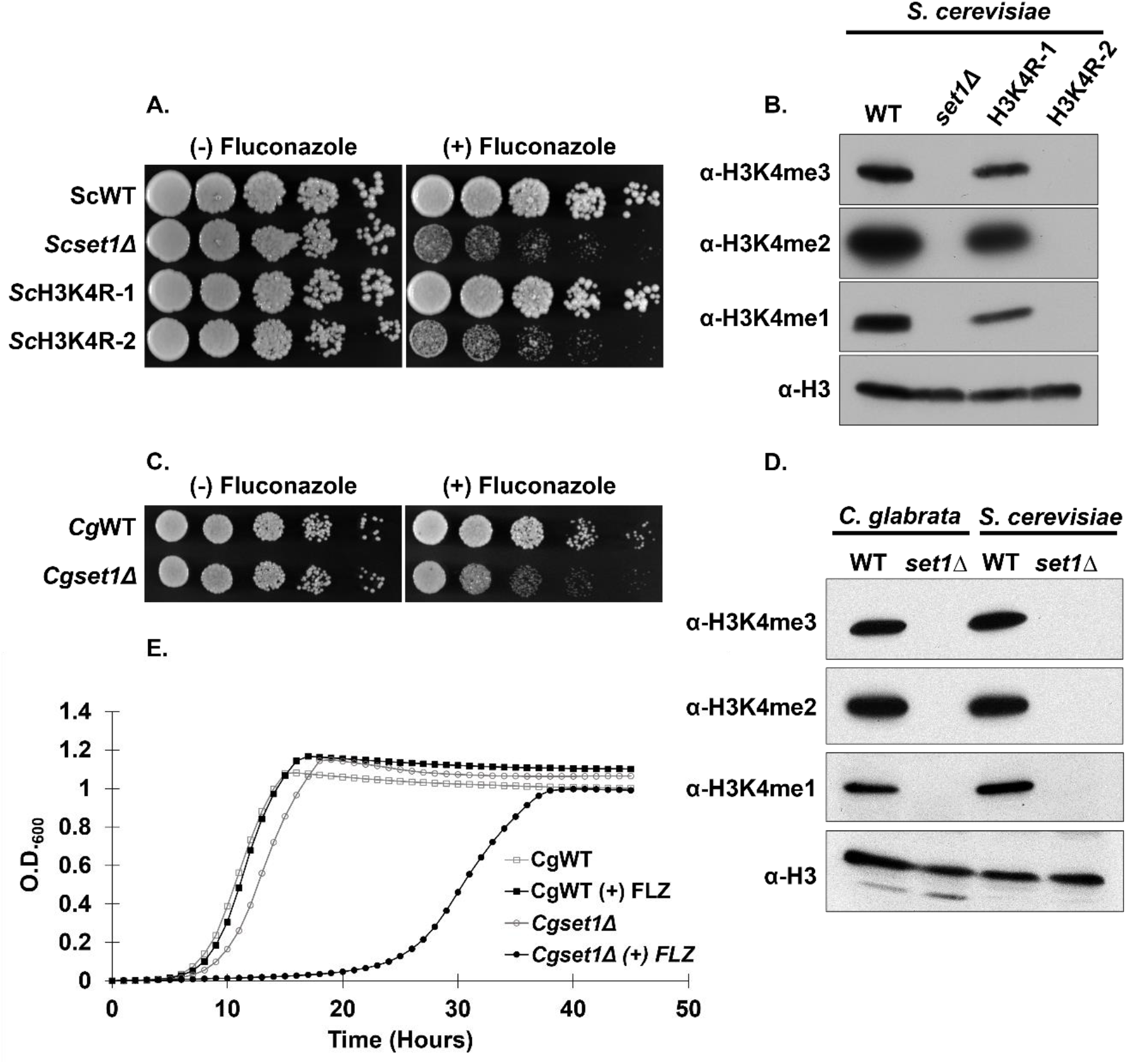
Loss of Set1-mediated mono-, di-, and trimethylation at histone H3K4 in Saccharomyces cerevisiae and Candida glabrata results in increased azole susceptibility and delayed growth in vitro. (A) Five-fold serial dilution spot assays of the indicated *S. cerevisiae* strains were grown on SC media with and without 8 µg/mL fluconazole and incubated at 30°C for 72 hours. (B & D) Whole cell extracts isolated from the indicated strains were immunoblotted using histone H3K4 methyl- specific mono-, di- and trimethylation antibodies of whole cell extracts isolated from the indicated strains. Histone H3 was used as a loading control. (C) Five-fold serial dilution spot assays of the indicated *C. glabrata* strains were grown on SC media with and without 32 µg/mL fluconazole and incubated at 30°C for 48 hours. (E) Liquid growth curve assay of the indicated *C. glabrata* strains grown over 50 hours with or without 32 µg/mL fluconazole.

To determine if an azole hypersensitive growth phenotype observed in *S. cerevisiae* is also conserved in the human fungal pathogen *C. glabrata*, WT (*Cg*WT) and a *set1Δ* (*Cgset1*Δ) strain were spotted on SC agar plates with and without 16 µg/mL fluconazole (Fig. 1C). Similar to what was observed in *S. cerevisiae*, deleting *SET1* in *C. glabrata* 2001 (CBS138, ATCC2001) showed an increase in azole susceptibility when compared to a *Cg*WT strain (Fig. 1C). Additionally, the *Cgset1Δ* strain had a significant growth delay in liquid growth cultures comparted to *Cg*WT when treated with 32 µg/mL fluconazole (Fig. 1E). Western blot analysis showed that deleting *CgSET1* abolished all histone H3K4 mono-, di-, and trimethylation confirming that *CgSET*1 is the sole histone H3K4 methyltransferase in *C. glabrata* (Fig. 1D). Altogether, our results show Set1-mediated histone H3K4 methylation in *S. cerevi*siae and *C. glabrata* is conserved and is necessary for maintaining a wild-type resistance to azole drugs.

### Loss of *C. glabrata* Set1 complex members alters azole efficacy and histone H3K4 methylation

In *S. cerevisiae*, Set1 forms a complex referred to as the <u>C</u>omplex <u>P</u>roteins <u>A</u>ssociated with <u>S</u>et1 or COMPASS. COMPASS forms a stable complex with 8 proteins which includes the catalytic subunit Set1, Swd1, Swd2, Swd3, Spp1, Bre2, Sdc1, and Shg1 (33–35). Previous studies in *S. cerevisiae* have determined that Swd1, Swd2, Swd3, Spp1, Bre2, and Sdc1 are necessary for Set1 to properly catalyze the various states of histone H3K4 mono-, di, and trimethylation (33–38) . To determine if COMPASS components are required to govern azole drug efficacy and Set1-mediated histone H3K4 methylation in *C. glabrata*, we generated deletion strains lacking *SET1*, *SPP1*, *BRE2* and *SWD1* and determined their MIC in RPMI media (Fig. 2A). Consistent with our agar and liquid growth assays in Figure 1, the *Cgset1Δ* strain showed increased susceptibility to fluconazole with an 8-fold difference in MIC compared to the *Cg*WT strain (Fig. 2A). A *Cgswd1Δ* strain showed a similar MIC as the *Cgset1Δ* strain while the MIC of *Cgspp1Δ* and *Cgbre2Δ* deletion strains were 4-fold different than the WT strain (Fig 2A). Furthermore, all *C. glabrata* COMPASS deletions strains showed an increase in susceptibility to azole drugs on agar plates similar to *S. cerevisiae* COMPASS deletion strains except for the *Scspp1Δ* which is likely due to differences in the histone H3K4 methylation status (Fig. 2B, 2C, S1A, and (29, 33, 34, 36, 38).

**FIG 2.**
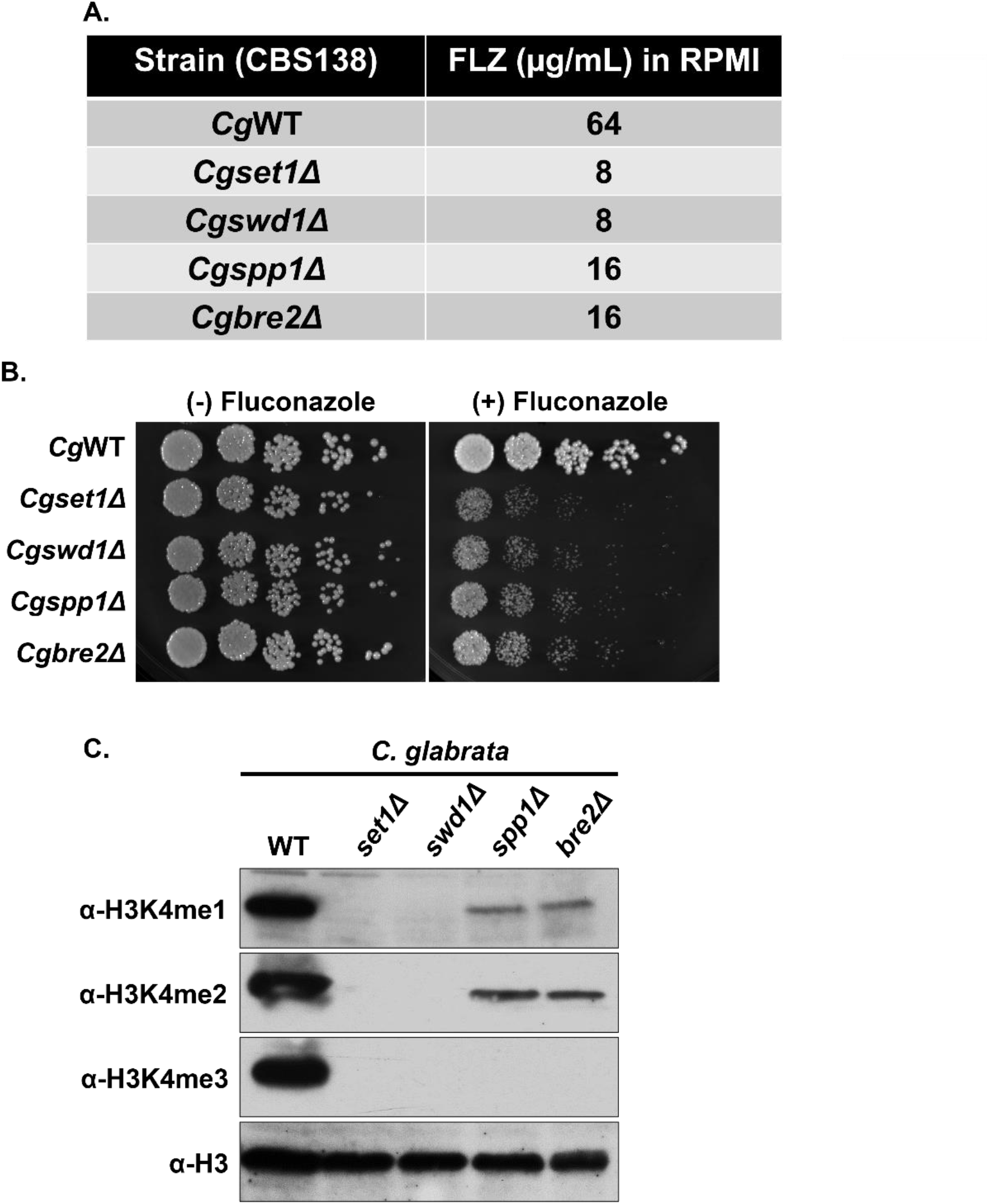
Deletion of Set1 complex members in C. glabrata results in increased azole susceptibility and loss of histone H3K4 methylation. (A) MIC assay of the indicated strains performed in RPMI 1640 media at 35°C and results recorded after 48 hours of incubation. (B) Five-fold serial dilution spot assays of the indicated *C. glabrata* strains were grown on SC plates with or without 32 µg/ml fluconazole. (C) Whole cell extracts isolated from the indicated strains were immunoblotted using H3K4 methyl-specific mono-, di- and trimethylation antibodies. Histone H3 was used as a loading control.

Western blot analysis determined that *Cgswd1Δ* strain lacked all forms of histone H3K4 methylation (Fig. 2C) which is also observed in *Cgset1Δ* and *Scset1Δ* strains (Fig. 2C and 1D). In contrast, deletion of *CgSPP1* and *CgBRE2* abolished all detectable levels of H3K4 trimethylation and significantly reduced the levels of histone H3K4 mono- and dimethylation. Taken together, our data show that when *C. glabrata* COMPASS subunits *SET1* and *SWD1* are deleted, global loss of histone H3K4 methylation is observed similar to what is seen when the subunits are deleted in *S. cerevisiae* (Fig 2C and (33, 34, 36, 38). However, the *Cgspp1Δ* has a total loss of histone H3K4 trimethylation and significant loss of histone H3K4 mono-and dimethylation similar to the *Cgbre2Δ* and *Scbre2Δ* strains (Fig 2C). For unknown reasons, the pattern of histone H3K4 methylation is different in the *Scspp1Δ* strain which only has a reduction in histone H3K4 trimethylation but not mono- or dimethylation (33–39). Altogether, these results suggest that the COMPASS complex is needed to mediate proper histone H3K4 methylation and WT resistance to azole drugs.

### The methyltransferase activity of Set1 governs azole drug efficacy in *C. glabrata*

To confirm that altered azole efficacy in the *Cgset1Δ* strain was due to loss of *SET1* and not a secondary mutation, a genomic fragment containing the *CgSET1* promoter, 5’UTR, open reading frame, and 3’UTR was amplified by PCR and cloned into the *C. glabrata* plasmid, pGRB2.0 (40). Because a H1017K mutation in the SET domain of *S. cerevisiae* Set1 is known to be catalytically inactive (28, 41, 42), we performed site-directed mutagenesis on pGRB2.0-*CgSET1* and generated an analogous mutation in *C. glabrata* Set1 at H1048K determined using the sequence alignment in Fig. 3A. Additionally, we deleted *SET1* in *C. glabrata* 2001HTU (ATCC200989) to utilize the *ura3* auxotrophic marker (43). Importantly, *Cg*2001HTU lacking *SET1* was hypersensitive to azole drugs similar to when *SET1* was deleted in *Cg*2001 (Fig. 1C and 3B).

**FIG 3.**
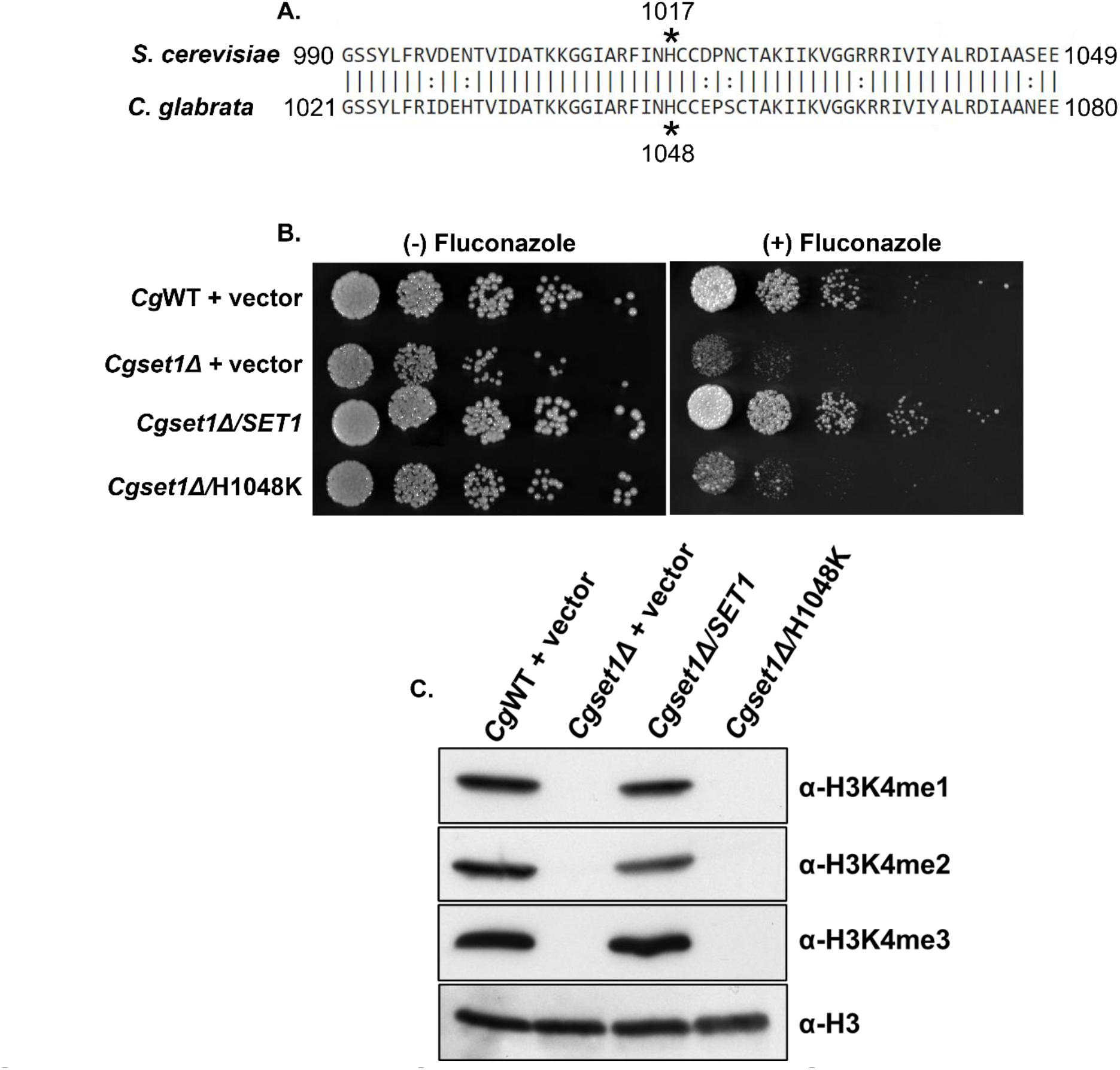
The catalytic activity of the SET domain is necessary for Set1-mediated histone H3K4 methylation and increased azole susceptibility in C. glabrata. (A) Five-fold serial dilution spot assays of the indicated C. glabrata strains were grown on SC plates with or without 32 µg/ml fluconazole. (B) Whole cell extracts isolated from the indicated strains were immunoblotted using methyl-specific mono-, di- and trimethylation antibodies. Histone H3 was used as a loading control.

Furthermore, transformation of pGRB2.0-*CgSET1* into the *Cg*2001HTU*/set1Δ* strain was able to rescue azole hypersensitivity while pGRB2.0-*Cgset1H1048K* did not rescue wild-type azole drug resistance as shown by serial dilution spot assays grown on SC agar plates with 32 µg/mL fluconazole (Fig. 3B). MIC assays under SC-ura conditions also show similar results (see Supplemental Fig. S1B). Western blot analysis indicated that pGRB2.0-*CgSET1* expression in *Cg*2001HTU*/set1Δ* strain restored histone H3K4 methylation to wild-type levels while *Cgset1H1048K* did not rescue histone H3K4 methylation confirming that this mutation lacks catalytic activity similar to *Scset1H1017K* (Fig. 3C). Importantly, quantitative real-time PCR analysis (qRT-PCR) confirmed that the plasmids expressing *CgSET1* and *Cgset1H1048K* were similar to the endogenously expressed *SET1* (Fig. S1C). This shows that loss of histone H3K4 methylation was not due to difference in expression levels but due to the catalytic inactivation of *Cgset1H1048K*. These data suggest that altered azole drug efficacy in *Cgset1Δ* strains are specifically due to the loss of *SET1* and its catalytic activity.

### Drug efflux pump expression and function is not altered in a *C. glabrata set1Δ* strain

In *Candida glabrata*, the major mechanisms for changes in drug resistance are due to changes in expression of *CDR1*, the main drug efflux pump, or gain-of-function mutations in *PDR1*, a gene that encodes the transcription factor for *CDR1* (7, 12, 19, 20, 44). To determine if altered drug resistance in *Cgset1Δ* cells was due to changes in *CDR1* or *PDR1* expression, we analyzed the transcript levels of *CDR1* and *PDR1* via qRT-PCR (Fig. 4A and B). We observed that *Cgset1Δ* cells grown with and without azoles do not significantly affect transcript levels of *CDR1* or *PDR1* when compared to a wild-type strain (Fig. 4A and B). Additionally, we analyzed the transcript levels of transporters *SNQ2, YOR1,* and *PDH1.* We did not see any significant changes in *SNQ2* or *YOR1,* but we did see a decrease in *PDH1* transcripts in a *set1Δ* strain upon azole treatment (Fig. S2). However, previous studies have shown loss of *PDH1* alone is not sufficient to lead to azole sensitivity (45). To determine if drug efflux was functional in *Cgset1Δ* cells, a Nile Red fluorescence-based assay was performed. Nile Red, a fluorescent lipophilic stain, has been shown to be a substrate for the ABC transporter Cdr1 in *C. albicans* and *C. glabrata* (46, 47). As a control, we also generated a *Cgpdr1Δ* strain, a deletion strain known to disrupt the expression of *CDR1* and subsequently prevent drug or Nile Red efflux (15). The Nile Red assay showed that *Cgset1Δ* cells had similar levels of Nile Red as wild-type cells but less Nile Red than *Cgpdr1Δ* cells (Fig. 4C). To induce *CDR1* expression levels, *Cgset1Δ* and wild-type cells were treated with fluconazole. Although azole treatment did reduce the amount of Nile Red in *Cgset1Δ* and wild-type cells compared to untreated cells, there was no discernable differences observed between *Cgset1Δ* and wild-type cells for their ability to efflux Nile Red, (Fig. 4C). Altogether these data suggest that cells lacking *SET1* have similar efflux capabilities as wild-type cells in the presence or absence of azole treatment. This suggests that the increase in azole sensitivity seen in a *Cgset1Δ* strain is not due to malfunction of Cdr1 expression or its efflux capabilities.

**FIG 4.**
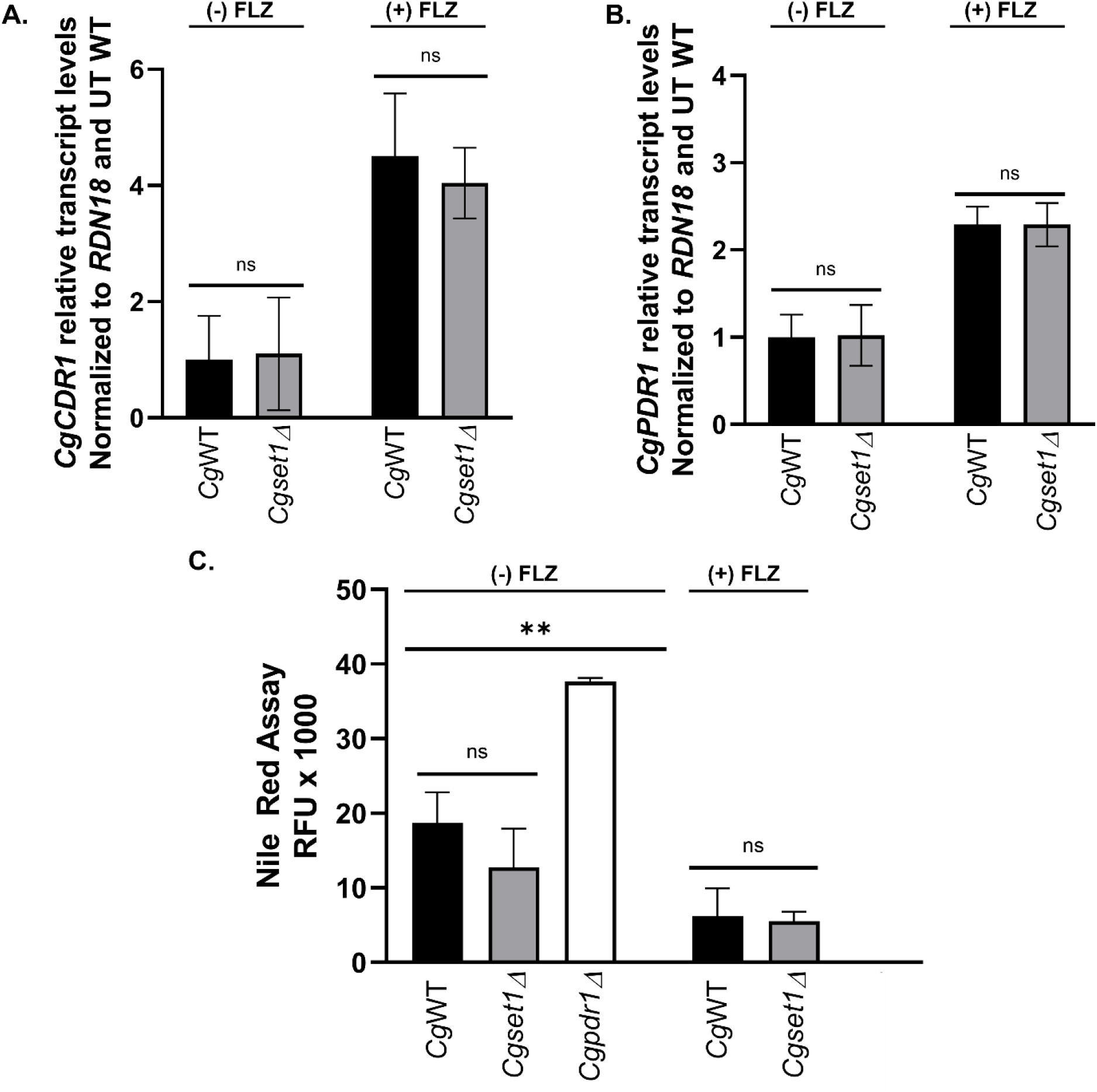
Deletion of SET1 in C. glabrata does not alter gene expression levels or function of the efflux drug transporter, CDR1 or transcription factor PDR1. (A and B) Expression of indicated genes was determined in *Cg*WT and *Cgset1Δ* strain cells treated with and without 64 µg/ml fluconazole for 3 hr by qRT-PCR analysis. Gene expression analysis was set relative to the untreated wild-type and expression was normalized to *RDN18* mRNA levels. Data were analyzed from ≥ 3 biological replicates with three technical replicates each. Error bars represent SD. (C) Red fluorescence units were measured as output in a Nile Red assay to determine the efficacy of Cdr1 in the indicated strains with and without fluconazole. A *pdr1Δ* strain was used as a control. Data were analyzed from ≥ 3 biological replicates with three technical replicates each. Statistics were performed using Graphpad Prism student t-test version 9.2.0. *ns represents p< 0.05,* \*\**p<0.01,* Error bars represent SD.

### Loss of Set1 leads to decreased expression of genes involved in the sterol biosynthesis pathway when treated with fluconazole

Because drug efflux function or transcript levels was not disrupted in a *Cgset1Δ* strain, we used RNA-sequencing analysis to provide insight into what gene pathway might be disrupted in the *Cgset1Δ* strain and explain why a loss of *SET1* alters azole drug efficacy. *Cg*WT and *Cgset1Δ* strains were treated with 64 µg/mL fluconazole for three hours in SC complete media and RNA was extracted for RNA-sequencing. Principle component analysis (PCA) and differentially expressed genes (DEG) analysis demonstrated by the volcano scatter plot (-log_2_ false discovery rate (FDR), y-axis) versus the fold change (x- axis) of the DEGs) indicate that the untreated and treated *Cg*WT strain is substantially and statistically different from the untreated and treated *Cgset1Δ* (Fig 5). DESeq2 analysis was used to identify the differentially expressed genes (DEGs) under fluconazole treatment using an FDR of 0.05. From this analysis, a total of 2389 genes were differentially expressed in *Cgset1Δ* vs. *Cg*WT under untreated condition (Fig. 5B). Whereas, 1508 genes were differentially expressed under treated conditions, where we observed 800 (14.2%) genes that were upregulated and 708 (12.6%) genes that were downregulated out of 5615 genes in *Cgset1Δ* compared to *Cg*WT (Fig. 5C and supplemental data). After applying a 1.4-fold cutoff to the data, we observed 1,644 genes differentially expressed in the untreated *Cgset1Δ* vs. *Cg*WT strains. In the treated strains, with a 1.4-fold cutoff, we observed 543 (9.7%) genes were down in a *Cgset1Δ* vs CgWT and 626 (11.1%) genes were up. These data show that *SET1* is important for maintaining proper gene expression in *C. glabrata*.

**FIG 5.**
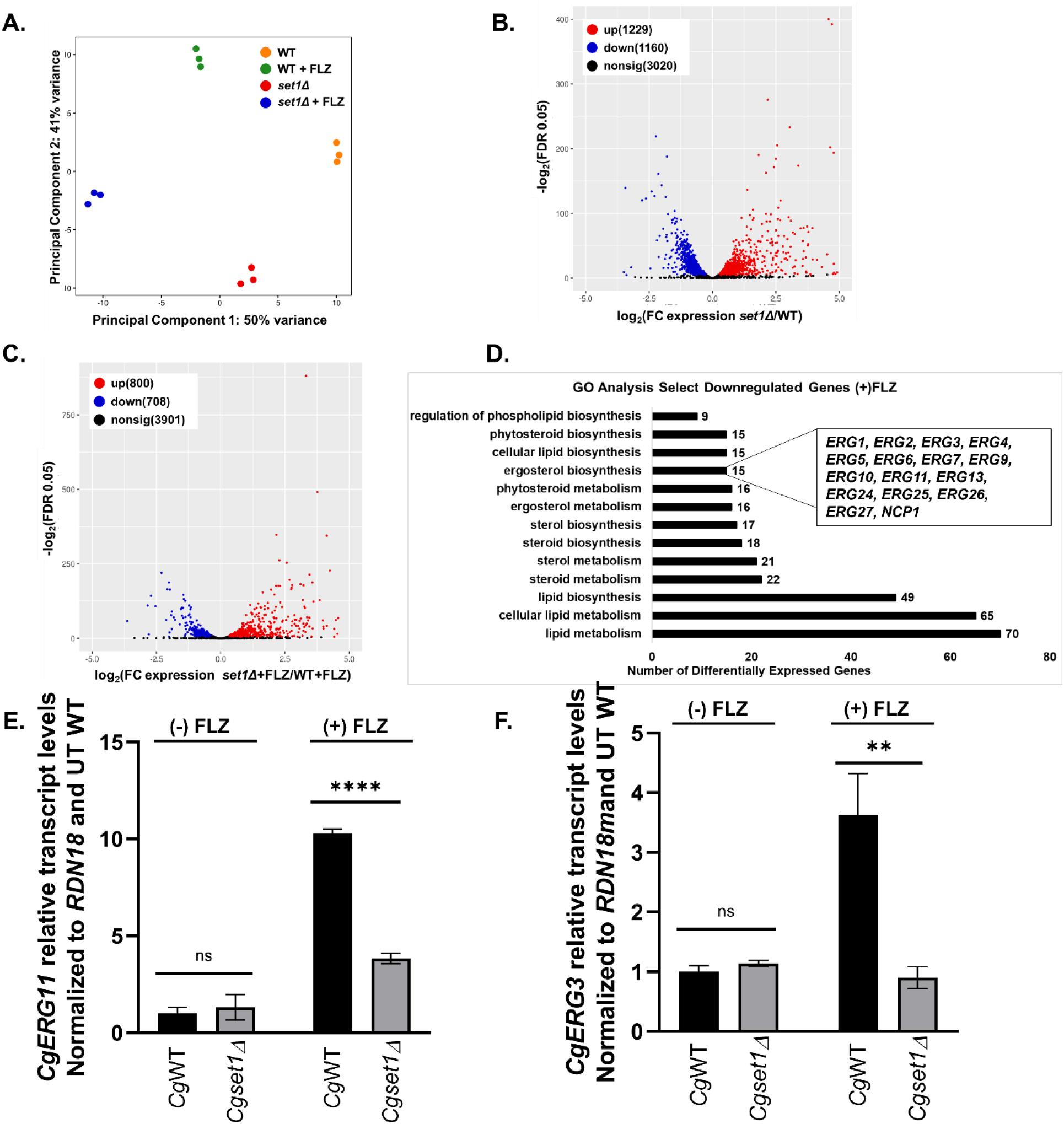
The deletion of SET1 in C. glabrata alters global and local levels of gene expression under untreated and azole conditions. The genome-wide changes in gene expression under azoles were performed using *C. glabrata* CBS138 WT and *set1Δ* strains. (A) The PCA for WT and *set1Δ* azole treated samples relative to WT untreated samples based on the counts per million. (B) Volcano plot showing the significance [−log_2_ (FDR), *y*-axis] *vs.* the fold change (*x*-axis) of the DEGs identified in the WT untreated samples relative to *set1Δ* untreated samples. (C) Volcano plot showing the significance [−log_2_ (FDR), *y*-axis] *vs.* the fold change (*x*-axis) of the DEGs identified in the *set1*Δ azole treated samples relative to WT azole treated samples. Genes with significant differential expression (FDR < 0.05) in (B and C) are highlighted in red or blue for up- and downregulated genes, respectively. Black highlighted genes are considered nonsignificant. (D) Genes from the RNA-seq dataset that were statistically significantly enriched (FDR < 0.05) were used for GO term determination of Set1- dependent DEGs under azole conditions. Downregulated genes refer to the DEGs that are dependent on Set1 for activation either directly or indirectly. Significantly enriched groups of GO terms were identified as the DEGs from only *set1*Δ and WT azole treated samples. (E and F) Expression of indicated genes was determined in WT and *set1Δ* strain cells treated with 64 µg/ml fluconazole for 3 hr by qRT-PCR analysis. Gene expression analysis was set relative to the untreated wild-type and expression was normalized to *RDN18* mRNA levels. Data were analyzed from ≥ 3 biological replicates with three technical replicates each. Statistics were performed using Graphpad Prism student t-test version 9.2.0.*\*\*\*\*p<0.0001 and **p=0.002.* Error bars represent SD.

Because Set1-mediated histone H3K4 methylation is known to play a key role in gene activation, we focused our attention on genes downregulated in *Cgset1Δ* compared to *Cg*WT. For azole-treated strains. GO Term Finder of the gene sets that were downregulated found significant GO terms involved in lipid, steroid and sterol/ergosterol metabolism or biosynthesis (Fig. 5D). For untreated strains, GO Term Finder identified significant GO terms involved in lipid metabolism but not steroid and sterol/ergosterol metabolism or biosynthesis (supplemental table S6). Interestingly, our data showed that 12 of the 12 genes involved in the late ergosterol biosynthesis pathway are down 1.4-fold or more in a *Cgset1Δ* compared to *Cg*WT under azole treated conditions (Fig. 5C, S4A & B and supplemental table S7). Whereas, 5 of the 12 late pathway *ERG* enzyme encoding genes were down in a *Cgset1Δ* compared to *Cg*WT under untreated conditions using a 1.4-fold difference in gene expression as a cutoff (Fig. 5D. and Supplemental table S7). Two of these differentially expressed genes *ERG11,* the gene that encodes the target of azoles, and *ERG3*, the gene that encodes the enzyme responsible for production of a toxic sterol when cells are treated with azoles, are known to play roles in azole drug resistance in various *Candida* species (17, 19, 48–50). To validate results seen in RNA-sequencing analysis, *ERG11* and *ERG3* transcript levels were analyzed by qRT-PCR. Our analysis showed that upon azole treatment, *ERG3* and *ERG11* transcript levels are induced in a WT strain (Fig. 5C and D) while loss of *SET1* prevented WT induction of both *ERG11* and *ERG3* under azole conditions. Even though our untreated RNA sequencing data set did show minor changes in *ERG3* and *ERG11* transcript levels, we did not detect any significant changes between *Cgset1Δ* and *Cg*WT cells when grown under untreated standard log-phase conditions using qRT-PCR analysis (Fig. 5E and F). We also performed gene expression analysis to determine if *ERG* gene transcript induction still depended on Set1 in saturated cultures. We show in both exponential and saturated cultures Set1 is necessary for *ERG3* and *ERG11* induction upon azole treatment in *C. glabrata* (Fig. 5E&F and S3A&B). Because *ERG3* transcript levels were decreased, we do not anticipate azole sensitivity is due to an increase in toxic sterols but by the lack of induction of *ERG11* and other *ERG* genes resulting in lower total cellular ergosterol levels (51, 52).

### Set1-mediated histone H3K4 methylation is enriched on *ERG* gene chromatin and is required for azole induction of *ERG* genes

Because histone H3K4 trimethylation is associated with gene induction, we wanted to determine if Set1 was directly catalyzing histone H3K4 methylation on chromatin at *ERG* loci. To determine if histone H3K4 trimethylation was present at *ERG11* and *ERG3* chromatin, chromatin immunoprecipitation (ChIP) analysis was performed using histone H3K4 trimethyl-specific antibodies. As expected, histone H3K4 trimethylation is highly enriched at the 5’-ends of the open reading frame of *ERG11* and *ERG3* in untreated conditions and further enriched upon azole treatment corresponding to increased transcript levels of *ERG11* and *ERG3* in both exponential and saturated cell cultures (Fig. 6A & B and S3D & E).

**FIG 6.**
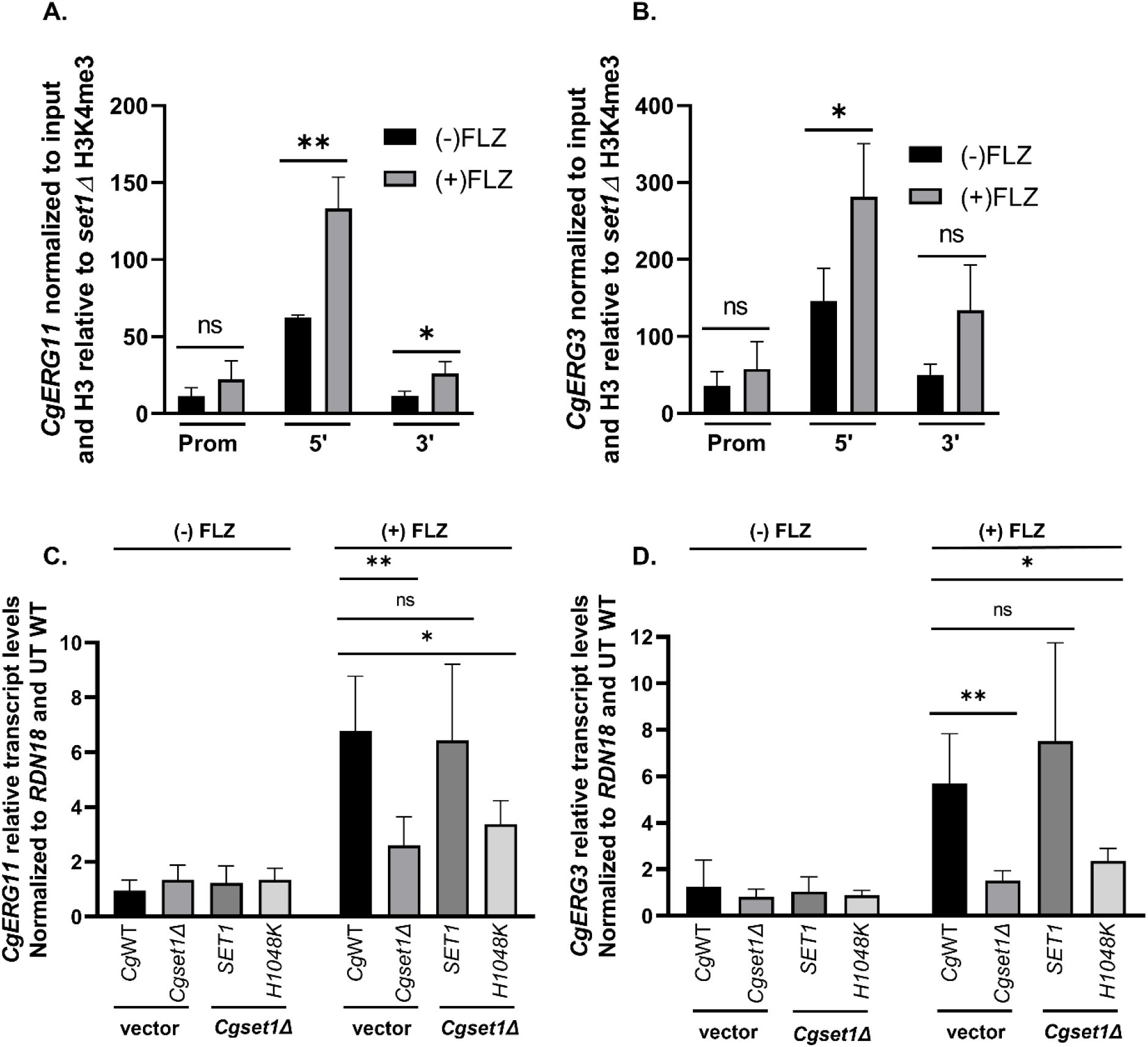
Histone H3K4 trimethylation is enriched on ERG gene chromatin and Set1- mediated histone H3K4 methyltransferase activity is required for azole induction of ERG genes. (A and B) ChIP analysis of histone H3K4 tri-methylation levels at the promoter, 5’, and 3’ regions of *ERG11* and *ERG3* in a wild-type *C. glabrata* strain with and without 64 µg/mL fluconazole treatment. ChIP analysis was set relative to a *set1Δ* strain and normalized to histone H3 and DNA input levels. Data were analyzed from 5 biological replicates with three technical replicates each, \**p<0.05.* (C and D) Expression of indicated genes was determined in the indicated mutants treated with and without 64 µg/ml fluconazole for 3 hr by qRT-PCR analysis. Gene expression analysis was set relative to the untreated wild-type containing an empty vector and expression was normalized to *RDN18* mRNA levels. Data were analyzed from ≥ 3 biological replicates with three technical replicates each. Statistics were performed using Graphpad Prism student t-test version 9.2.0. *\*\*\*\*p<0.0001 and **p<0.01.* Error bars represent SD.

To confirm that this was due to the methyltransferase activity of Set1, we performed qRT-PCR transcript analysis using the *Cg*2001HTU*set1Δ* strain expressing pGRB2.0 only, pGRB2.0-*CgSET1*, and pGRB2.0-*Cgset1H1048K*. *Cg*2001HTU expressing pGRB2.0 only was used as our WT control. As shown in Figure 6C and D, pGRB2.0-*CgSET1* was able to induce *ERG11* and *ERG3* similar to WT cells under azole treatment indicating that *SET1* expression could rescue the *ERG* gene expression in the *Cg*2001HTU*set1Δ* strain. This rescue of *ERG* gene expression was dependent on the catalytic activity of Set1 since expression of pGRB2.0-*Cgset1H1048K* did not restore *ERG* gene expression under azole treatment. Additionally, it looked similar to the *Cg*2001HTU*set1Δ* strain expressing pGRB2.0 indicating that the catalytic activity of Set1 is required for azole gene induction. Altogether, these data show that Set1- mediated histone H3K4 methylation directly targets the chromatin of *ERG* genes, and this epigenetic modification is required for azole induction of *ERG* genes. Based on our results, the lack of Set1 or histone H3K4 methylation on *ERG11* chromatin prevents the transcriptional response for inducing *ERG* genes which consequently disrupts ergosterol homeostasis, thus making the *Cgset1Δ* strains more susceptible to azole drugs.

## DISCUSSION

In this study, we established that loss of Set1-mediated histone H3K4 methylation alters azole drug susceptibility in *S. cerevisiae* and *C. glabrata*. This increase in susceptibility to azole drugs in a *Cgset1Δ* strain was not because of the typical changes in *CDR1* and *PDR1* expression levels or their ability to efflux drugs. However, we observed that strains lacking histone H3K4 methylation failed to induce *ERG* genes. This azole-induced gene expression was dependent on Set1 methyltransferase activity and correlated with gene-specific increases in histone H3K4 trimethylation on chromatin at *ERG* genes (see model, Fig. 7). Overall, we have provided an epigenetic mechanism upon azole treatment that is dependent on histone H3K4 methylation governing ergosterol homeostasis. Identifying and understanding this Set1-*ERG* pathway and other epigenetic mechanisms contributing to altered drug susceptibility will be important for the development of alternative drug targets that could be used in combinatorial therapy for treating patients with drug resistant fungal infections.

**FIG 7.**
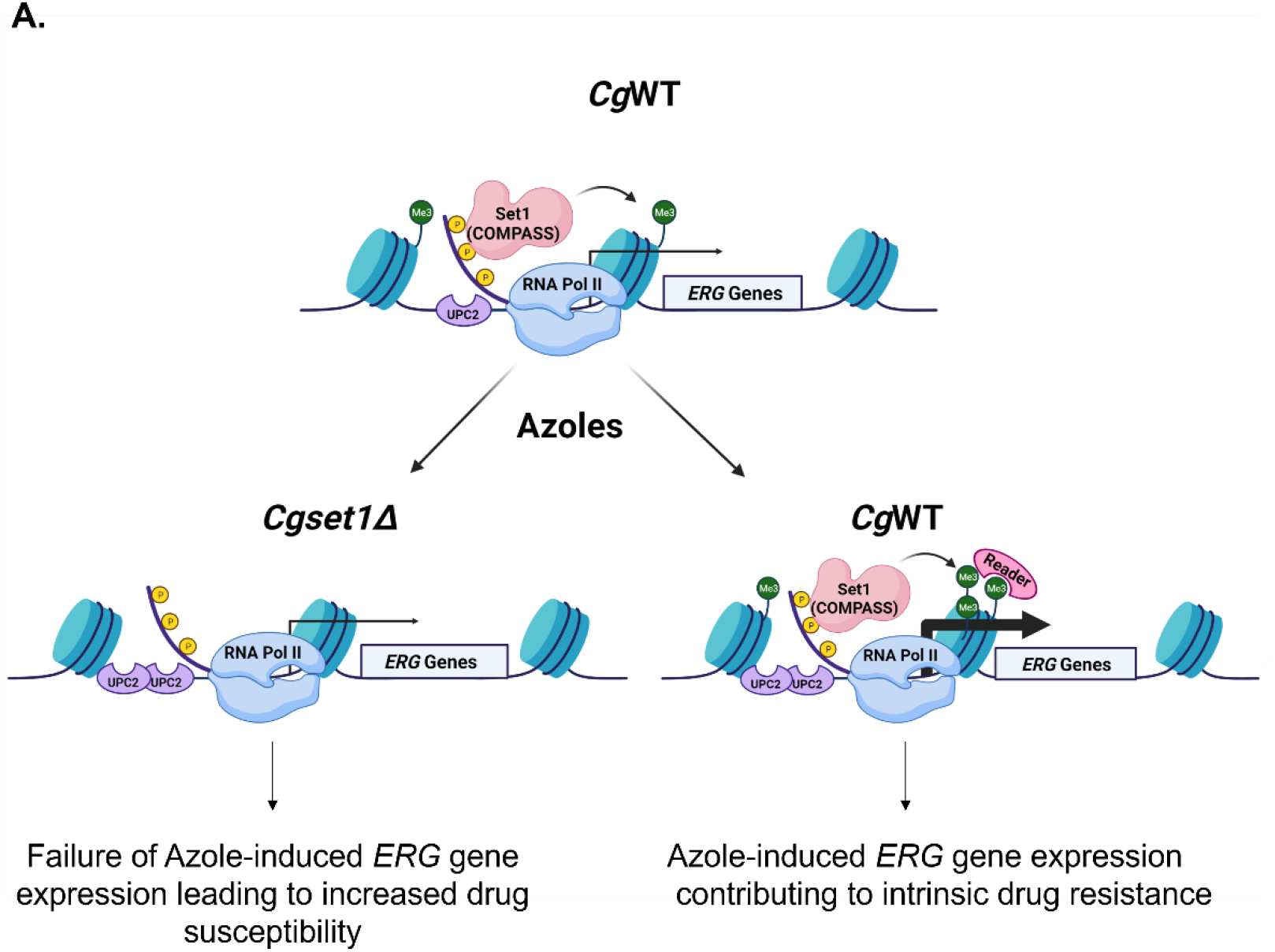
Model for the role of Set1-H3K4 methylation in epigenetic control of ERG genes (Biorender). (A) Under aerobic conditions, the Set1 complex mediates histone H3K4 methylation on chromatin at *ERG* genes. In the presence of azoles, azole-induced transcriptional activation recruits TFs, RNA Polymerase II, and the Set1 complex to increases Histone H3K4 methylation. This increase in methylation could permit additional recruitment of other co-factors/epigenetic regulator (e.g., Set3 and/or SAGA complex) that contain “reader” domains that recognize and bind to the H3K4 methyl mark. Thus, this Set1-Erg pathway contributes to the intrinsic azole resistance in *C. glabrata.* In the absence of Set1, histone H3K4 methylation is abolished and failure of recruiting additional H3K4 methyl “readers” prevent the induction of *ERG* genes, thus making the *C. glabrata* more susceptible to azole treatment.

Set1 is the catalytic subunit of a multi-subunit protein complex called COMPASS that mono-, di-, and trimethylates histone H3K4. In this study, we show that *C. glabrata* Set1 is the sole histone H3K4 methyltransferase under log-phase growth conditions since deletion of *SET1* abolishes all forms of histone H3K4 methylation similar to what is seen in *S. cerevisiae* and *C. albicans*. Deletion of the genes encoding *C. glabrata* COMPASS complex subunits Swd1 and Bre2 have similar loss of histone H3K4 methylation as their *S. cerevisiae* counterparts (see Fig 2C and (33, 34, 36, 38)). However, deleting *SPP1* in *C. glabrata* abolishes all histone H3K4 trimethylation and significantly reduces the levels of histone H3K4 mono- and dimethylation which is in contrast to what is found in a *Scspp1Δ* strain where only histone H3K4 trimethylation is disrupted but retains WT levels of mono- and dimethylation (33, 34, 36, 38). We speculate that this difference in histone H3K4 methylation pattern is due to how *Cg*Spp1 assembles with the COMPASS complex allowing *Cg*Spp1 to have a greater impact on the overall catalytic activity of COMPASS. Interestingly, this pattern of methylation appears to correlate with sensitivity to azole drugs (compare Fig. 2B with supplemental Fig. S1A) where *Scspp1Δ* grows more similar to a WT strain than *Cgspp1Δ* when grown on azole containing plates.

Our published observation and current data show that loss of *SET1* alters ergosterol homeostasis and azole susceptibility in *S. cerevisiae* and *C. glabrata* (28, 29). Specifically, loss of *SET1* in *S. cerevisiae* altered expression of genes involved in ergosterol biosynthesis under untreated conditions (28). In contrast, in *C. glabrata*, significant changes in *ERG* gene expression were only observed under azole treated conditions but not untreated conditions suggesting that histone H3K4 methylation is needed for azole induced gene induction and not basal level expression (Fig. 5E and F). Although regulation of ergosterol biosynthesis has been shown to be coupled to expression of ABC transport genes such as *CDR1* and its transcription factor Pdr1 (53, 54), compensatory changes in *CDR1 and PDR1* expression levels was not observed in a *cgset1Δ* strain when treated with azole drugs (Fig. 4A and 4B). More investigation will be needed to understand how *CDR1* and/or *PDR1* are epigenetically regulated in *C. glabrata* (26, 55).

In *C. albicans*, Raman et al., reported that loss of *SET1* in did not alter azole sensitivity but did decrease virulence in mice (30). Furthermore, *C. albicans* is naturally more susceptible to azole drugs than *S. cerevisiae* and *C. glabrata*. We suspect the difference observed in these organisms in azole sensitivity and gene regulation is likely due to their differences in sterol uptake. For example, *C. glabrata and S. cerevisiae* can uptake sterols under a variety of conditions where *C. albicans* does not (56). Interestingly, loss of *SET1* in *S. cerevisiae* can also permit sterol uptake under aerobic conditions since sterol transporter transcripts of *PDR11* and *AUS1* are increased in a *Scset1Δ* strain (28). In contrast, *AUS1,* is constitutively expressed in *C. glabrata* under aerobic and anaerobic conditions allowing sterol uptake. Even though *AUS1* is constitutively expressed, we do observe a slight increase in *AUS1* transcript levels in a *Cgset1Δ* relative to *Cg*WT under untreated conditions but not treated conditions (see supplemental data Fig. S3C). Alternatively, loss of *SET1* may not alter azole efficacy in *C. albicans* because it does not regulate the expression of *ERG* genes or sterol transporters. However, other epigenetic factors as indicated below are likely playing a role in *C. albicans*. Nonetheless, additional studies will be needed to determine the precise mechanistic cause of these distinct differences.

Overall, our data suggest that histone H3K4 methylation is an epigenetic mechanism to help induce *ERG* gene expression when *C. glabrata* strains are exposed to azole drugs. We propose histone H3K4 methylation and possibly other epigenetic marks are contributing factors to *C. glabrata’s* natural resistance to azole drugs. Interestingly, several histone deacetylases (HDACs) have been implemented in azole resistance in *C. albicans* such as *Ca*Hda1, *Ca*Rpd3, and *Ca*Hos2 (22, 23, 57–59). Additionally, HDAC inhibitors have been shown to have a synergistic effect on cells when combined with azoles and echinocandins (57, 58, 60, 61). Interestingly, the treatment of *C. albicans* with trichostatin A (TSA) lacks the trailing effect observed in MIC assays when using azole drugs and the lack of trailing effect was attributed to reduced *CDR* and *ERG* gene expression (58, 62). In a similar manner, *Cgset1Δ* also lacks a trailing effect in our MIC assays (personal observation) which we suspect is specifically due to the lack of azole-induced *ERG* gene expression since *CDR1* expression was not altered (Fig. 4A and 5). Furthermore, treatment of drug resistant fungal pathogens including various isolates of *C. glabrata* with a Hos2 inhibitor MGCD290 showed synergy with azole drugs which converted the MICs of azole treatment from resistant to susceptible (60). Since Hos2 is known to be a key component of the Set3 complex and the Set3 complex is recruited to chromatin via Set1-mediated histone H3K4 methylation (63, 64), it is likely MGCD290 is mediating its effect with azoles through inhibiting azole-induced *ERG* gene expression.

We expect that the Set1-*ERG* regulatory pathway controlling ergosterol homeostasis will not only impact drug resistance but will also impact fungal pathogenesis. For example, *C. albicans* strains lacking *ERG11* or *ERG3* produces avirulent hyphae, decreases the adherence to epithelial cells, and reduces virulence of *C. albicans* in oral mucosal infections and disseminated candidiasis (65–67). Similarly, deletion of *SET1* in *C. albicans* also forms hyphae, decreases epithelial adherence, and reduces virulence of *C. albicans* in disseminated candidiasis (30). Based on our current data in *C. glabrata*, we speculate that loss of *SET1* in *C. albicans* reduces expression of *ERG* genes and ergosterol production which in turn reduces epithelial adherence and thus alters the virulence of *C. albicans*. Therefore, the loss of *SET1* could also alter the virulence of *C. glabrata*. Interestingly, several genes encoding cell wall proteins and adhesion factors are also down regulated in a *Cgset1Δ* strain as determined by RNA-sequencing. However, future studies will be needed to determine if this Set1-*ERG* regulatory pathway exists for *C. albicans* and if *SET1* is controlling virulence factors for *C. glabrata*.

Overall, the occurrence of multidrug resistant strains is increasing across all *Candida* species. In addition, with the development and identification of multidrug resistant fungal species such as *C. auris*, a pathogen of urgent concern for the CDC, it is imperative to find alternative treatment options. Our study along with others provide compelling evidence that epigenetic modifiers are playing key roles in fungal pathogenesis and drug resistance. Understanding these epigenetic events and the pathways they impact are needed to develop new drug therapies so that current and newly emerging multidrug resistant fungal pathogens can be effectively treated.

## MATERIALS AND METHODS

### Plasmids and yeast strains

All plasmids and yeast strains are described in Table S1 and S2. *C. glabrata* 2001 (CBS138, ATCC2001) and *C. glabrata* 2001HTU (ATCC200989) were purchased from ATCC (43). A genomic fragment containing the *CgSET1* promoter, 5’UTR, open reading frame, and 3’UTR was amplified by PCR and cloned into the pGRB2.0 plasmid. The pGRB2.0 plasmid was purchased from Addgene. Standard, site-directed mutagenesis was used to generate *Cgset1H1048K*. *Candida glabrata SET1*, *BRE2*, *SWD1*, *SPP1*, and *PDR1* genes were deleted via standard homologous recombination. Briefly, drug resistant selection markers were PCR amplified with Ultramer DNA Oligos (IDT) using pAG32-HPHMX6 (hygromycin) or pAG25-NATMX6 (nourseothricin).

### Serial dilution spot and liquid growth assays

For serial dilution spot assays, yeast strains were inoculated in SC media and grown to saturation overnight. Yeast strains were diluted to an OD_600_ of 0.1 and grown in SC media to log phase shaking at 30°. The indicated strains were spotted in five-fold dilutions starting at an OD_600_ of 0.01 on untreated SC plates or plates containing 16,32, or 64 µg/ml fluconazole (Sigma-Aldrich, St. Louis, MO). Plates were grown at 30° for 1-3 days. For growth assays, the indicated yeast strains were inoculated in SC media and grown to saturation overnight. Yeast strains were diluted to an OD_600_ of 0.1 and grown in SC media to log phase shaking at 30°. The indicated strains were diluted to an OD_600_ of 0.0001 in 100 μl SC media. Cells were left untreated or treated with 64 μg/ml fluconazole and grown for 50 hrs shaking at 30°. The cell density OD_600_ was determined every 1 hr using the Bio-Tek Synergy 4 multimode plate reader.

### Cell extract and Western blot analysis

Whole cell extraction and western blot analysis to detect histone modifications were performed as previously described (36, 68). The histone H3K4 methylation-specific antibodies were used as previously described; H3K4me1(Upstate 07-436, 1:2,500), H3K4me2 (Upstate 07-030, 1:10,000), H3K4me3 (Active motif 39159, 1:100,000) (28, 69). Histone H3 antibodies were used as our loading control (Abcam ab1791, 1:10,000).

### RNA-sequencing analysis

The CBS138 *Cg*2001 WT and *set1*Δ strains were inoculated in SC media and grown to saturation overnight. Cells were diluted to an OD_600_ of 0.1 and recovered to log phase for 3 hours shaking at 30°. Prior to treatment, cells were collected for the untreated sample and zero time point. Cultures were treated at an OD_600_ of 0.2 with 64 µg/ml fluconazole (Sigma-Aldrich, St. Louis, MO) dissolved in DMSO as previously described (70). Cells were collected after 3 hours. Total RNA of three biological replicates for each condition and sample were isolated by standard acid phenol purification, treated with DNase (Ambion), and total RNA was purified using standard acid phenol purification. The quality of the RNA was tested using an Agilent Bioanalyzer 2100 using the High Sensitivity DNA Chip. The complementary DNA library was prepared by the Purdue Genomics Facility using the TruSeq Stranded Kit with poly(A) selection (Illumina) according to the manufacturer’s instructions. The software Trimmomatic v.0.39 was used to trim reads, removing adapters and low quality bases (71). STAR v.2.5.4b was used to align reads to the *C. glabrata* CBS 138 reference genome, version s02-m07-r23 (72). One mismatch was allowed per read. HTSeq v.0.6.1 was used to generate the gene count matrix on “intersection-nonempty” mode (73). R version 3.5.1 and Bioconductor release 3.6 were used to perform all statistical analyses on the RNA-seq data. The intersection of genes identified as statistically significantly differentially expressed with a Benjamini-Hochberg corrected false discovery rate of less than 5% by DESeq2 v.1.18.0 was used in downstream analyses (74, 75).

### Quantitative real-time PCR analysis

RNA was isolated from cells by standard acid phenol purification. Reverse transcription was performed using the ABM all-in-one 5X RT Mastermix kit (ABM, Richmond, Canada). Primer Express 3.0 software was used for designing primers and quantitative real-time polymerase chain reaction (qRT-PCR) was performed as previously described (28, 76, 77). A minimum of three biological replicates, including three technical replicates, were performed for all samples. Data were analyzed using the ΔΔCt method where *RDN18* (18S ribosomal RNA) was used as an internal control. All samples were normalized to an untreated, untagged WT strain.

### Minimal inhibitory concentration assay

MIC assays were performed based on a modified version of the CLSI method for testing yeast, 3^rd^ addition (78). Briefly, yeast strains were inoculated in SC media and grown to saturation overnight. The indicated strains were diluted to an OD_600_ of 0.003 in in SC or RPMI media. Cells were mixed with fluconazole (Cayman Chemical) for a final volume of 100µl per well in a 96 well polystyrene microplate with concentrations of fluconazole ranging from 0-256 µg/mL. Plates were incubated at 35°C and MICs were recorded at 24 hours. MICs were determined visually and were counted as wells where >90% of growth was inhibited.

### Nile Red Assay

Fluorescence-based Nile red assays were performed as previously described (46). Briefly, cells were grown overnight in SC media to saturation. Cells were back diluted to an O.D._600_ of 0.1 and grown for 6 hours. Cells were collected then washed with PBS twice and incubated in 1.5mL of PBS+2% glucose for 1 hour. Next, 2.87 µL of a 1mg/mL stock of Nile red (Sigma) was added to each sample and incubated at 30°C shaking for an additional 30 minutes. Samples were washed twice with PBS and placed in triplicate in a black 96-well flat-bottomed polystyrene microplate. Fluorescence was detected using a Bio-Tek Synergy 4 multimode plate reader using an excitation wavelength of 553nm and an emission wavelength of 636nm.

### Chromatin Immunoprecipitation

ZipChIP was performed as previously described (69). Briefly, 50 ml cultures were grown to log phase (OD_600_ of 0.6) in SC complete media at 30° shaking. Treated cells were dosed with 64µg/mL fluconazole (Cayman Chemical) at an OD_600_ of 0.2 for 3 hours. Additionally, cultures were grown to saturation, back diluted to an O.D._600_ of 0.6, treated with 64µg/mL fluconazole for 3 hours and collected. Cells were formaldehyde cross- linked and harvested as previously described (69). Cell lysates were precleared with 5 µl of unbound Protein G magnetic beads for 30 min rotating at 4°. A total of 12.5 µl of precleared lysate was immunoprecipitated with 10 µl of Protein G magnetic beads (10004D; Life Technologies) conjugated to 1 µl of Histone H3K4me3 antibody (Millipore 07-473) or Histone H3 antibody (Abcam ab1791). Probe sets used in qRT-PCR are described in supplemental Table S5.

## ACKNOWLEGEMENTS

This work was supported by grants from the National Institutes of Health to S.D.B (AI136995), the Purdue Department of Biochemistry Bird Stair Fellowship (to K.M.B.), Purdue Center for Cancer Research (Grant P30CA023168: DNA Sequencing Shared Resource and Collaborative Core for Cancer Bioinformatics at Purdue), Walther Cancer Foundation and the IU Simon Cancer Center (Grant P30CA082709). Additional funding support was provided by the NIFA 1007570 (S.D.B). We thank the Purdue Bioinformatics Core for their pipelines and bioinformatics software.

---

**FIG S1.**
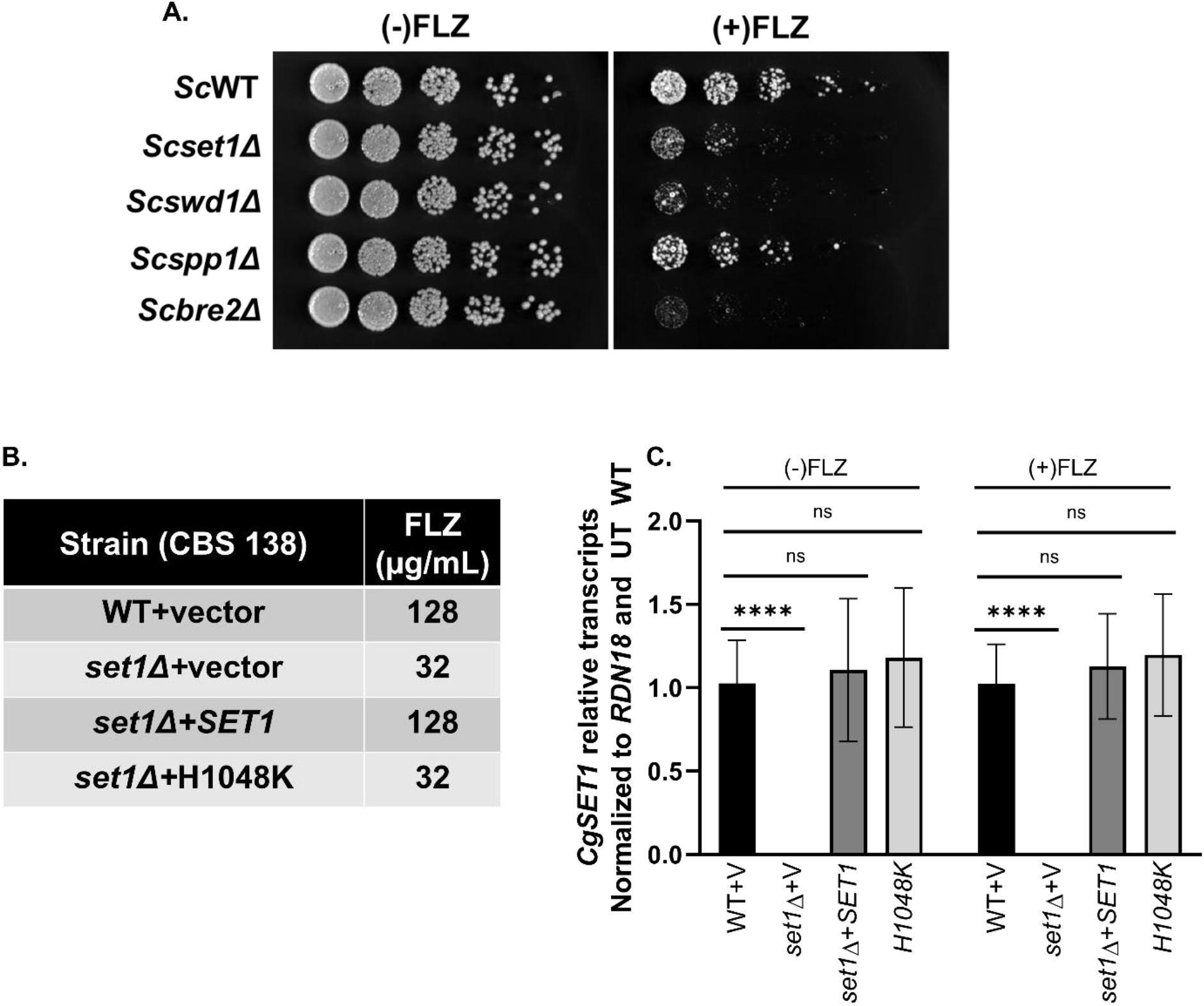
Loss of Set1 complex members results in altered azole efficacy. (A) Five-fold serial dilution spot assays of the indicated S. cerevisiae strains were grown on SC plates with or without 8 µg/ml fluconazole. (B) MIC assay of the indicated strains performed in SC media at 35°C and results recorded after 24 hours of incubation. (C) Expression of SET1 was determined in the indicated mutants treated with and without 64 µg/ml fluconazole for 3 hr by qRT-PCR analysis. Gene expression analysis was set relative to the untreated wild-type and expression was normalized to RDN18 mRNA levels. Data were analyzed from 4 biological replicates with three technical replicates each. Statistics were performed using Graphpad Prism student t-test version 9.2.0. ****p<0.0001. Error bars represent SD.

**Fig S2.**
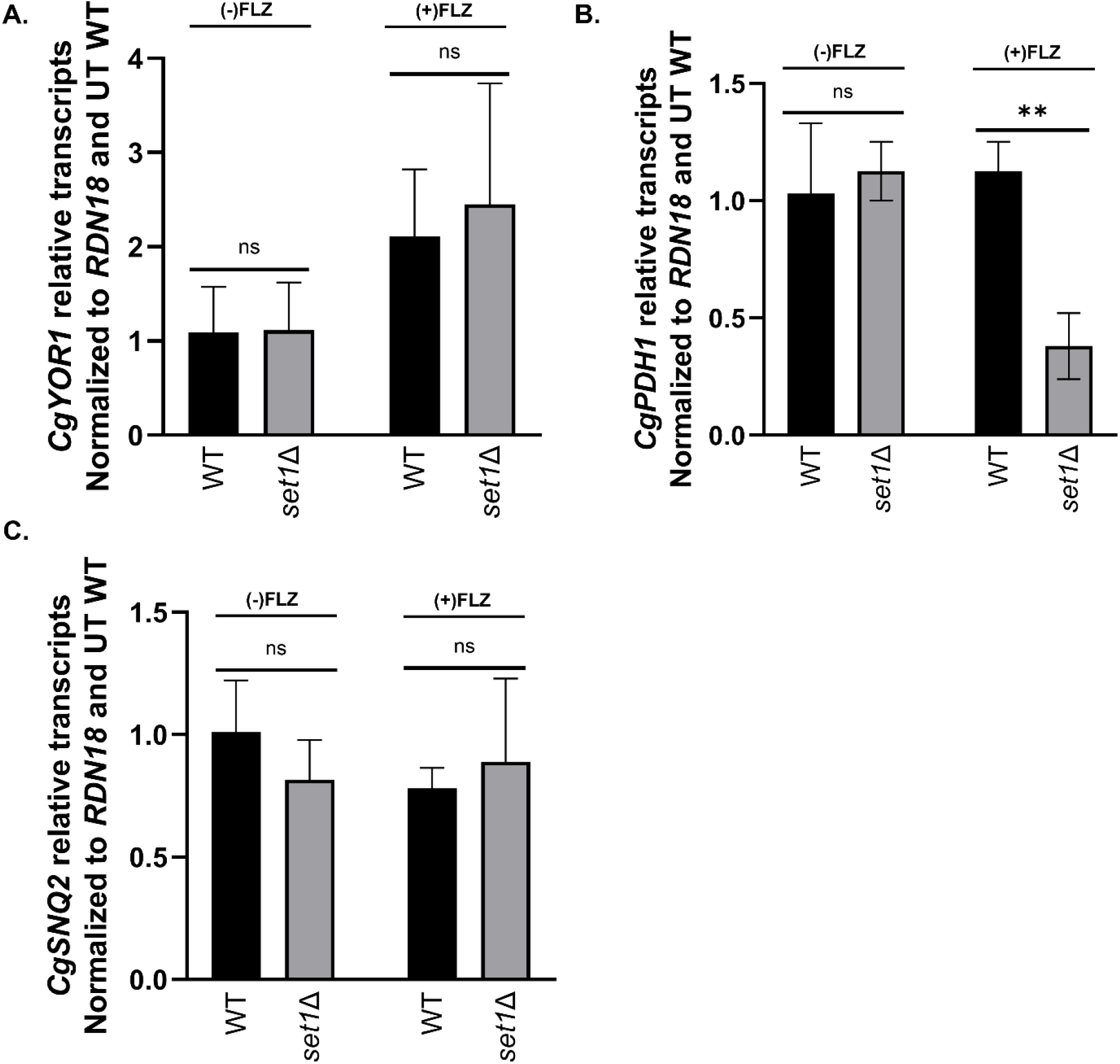
Transcript levels of drug transporters in a set1Δ strain compared to wild-type. Expression of the indicated genes were determined in the indicated mutants treated with and without 64 µg/ml fluconazole for 3 hr by qRT-PCR analysis. Gene expression analysis was set relative to the untreated wild-type and expression was normalized to *RDN18* mRNA levels. Data were analyzed from ≥ 3 biological replicates with three technical replicates each. Statistics were performed using Graphpad Prism student t-test version 9.2.0. *\*\*p<0.01.* Error bars represent SD.

**Fig S3.**
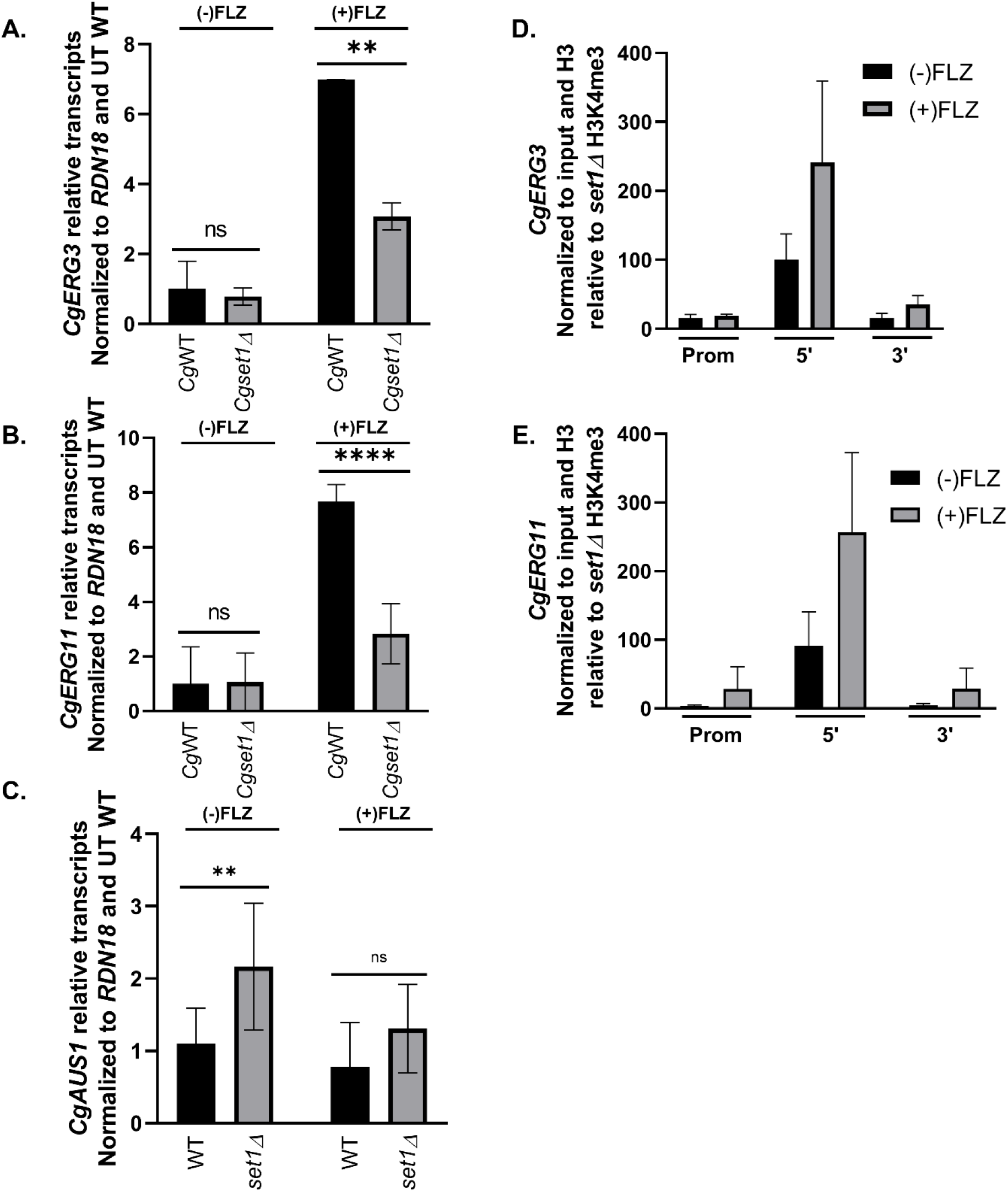
Histone H3K4 trimethylation is enriched on ERG gene chromatin and Set1- mediated histone H3K4 methyltransferase activity is required for azole induction of ERG genes in saturated cells. (A and B) Expression of genes was determined in the indicated strains treated with and without 64 µg/ml fluconazole in a saturated culture for 3 hr by qRT-PCR analysis. Gene expression analysis was set relative to the untreated wild-type expression was normalized to *RDN18* mRNA levels. Data were analyzed from ≥ 3 biological replicates with three technical replicates each. (C) Expression of genes was determined in the indicated strains treated with and without 64 µg/ml fluconazole in an exponential culture for 3 hr by qRT-PCR analysis. Gene expression analysis was set relative to the untreated wild-type expression was normalized to *RDN18* mRNA levels. Data were analyzed from ≥ 3 biological replicates with three technical replicates each. (D and E) ChIP analysis of histone H3K4 tri-methylation levels at the promoter, 5’, and 3’ regions of *ERG11* and *ERG3* in a wild-type *C. glabrata* strain with and without 64 µg/mL fluconazole treatment in saturated cell cultures. ChIP analysis was set relative to a *set1Δ* strain and normalized to histone H3 and DNA input levels. Data were analyzed from 3 biological replicates with three technical replicates each. Statistics were performed using Graphpad Prism student t-test version 9.2.0. \**p<0.05. ****p<0.0001 and **p<0.01.* Error bars represent SD.

**Fig S4.**
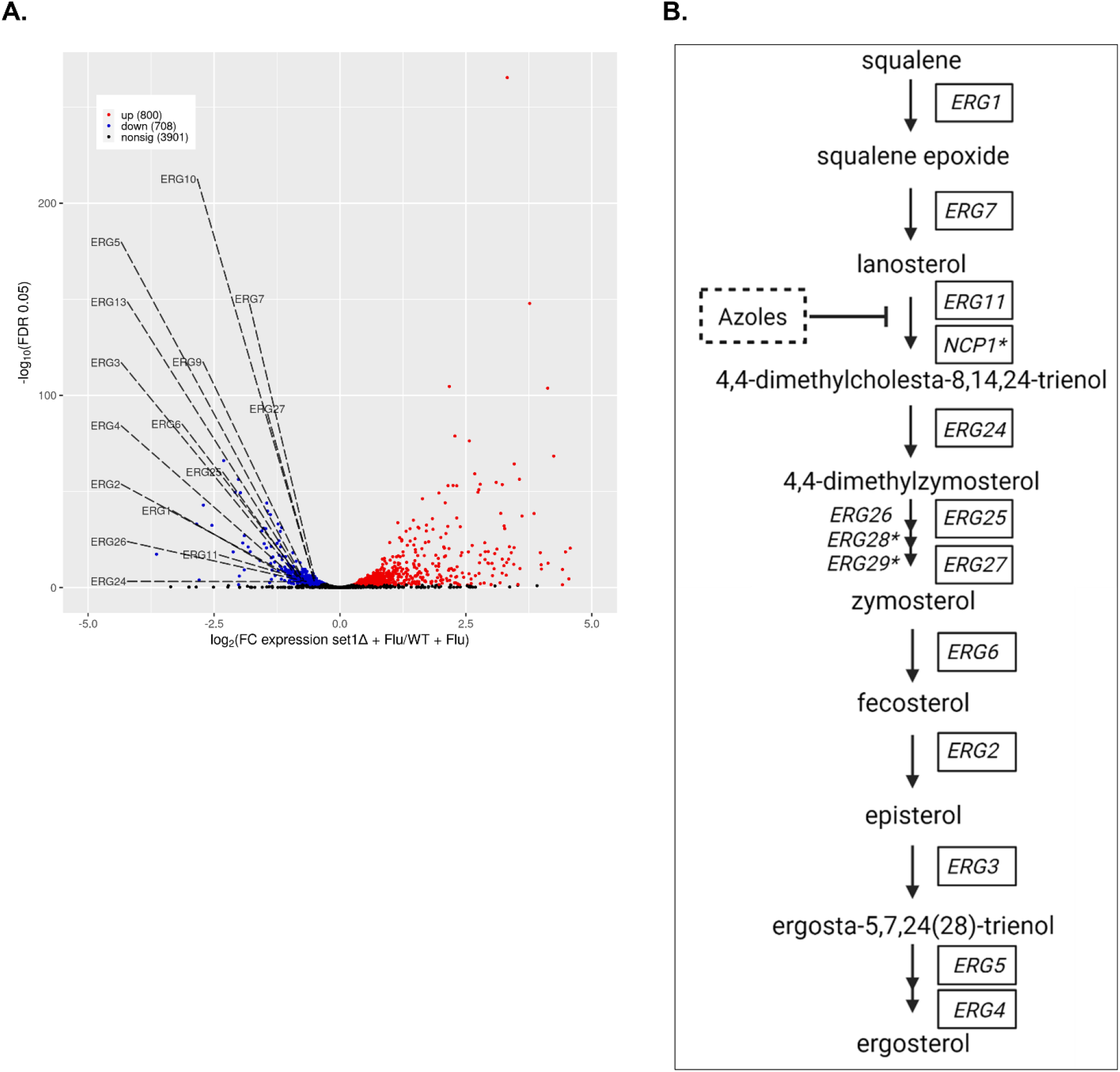
Genes encoding enzymes of the late ergosterol pathway are down in a set1Δ strain upon azole treatment in C. glabrata. (A) Volcano plot showing the significance [−log_2_ (FDR), *y*-axis] *vs.* the fold change (*x*-axis) of the DEGs identified in the *set1*Δ azole treated samples relative to WT azole treated samples. Genes with significant differential expression (FDR < 0.05) are highlighted in red or blue for up- and downregulated genes, respectively. Down-regulated *ERG* genes are labelled in the plot which include 12 of the *ERG* genes in the late pathway and two *ERG* genes in the early pathway. Black highlighted genes are considered nonsignificant. (B) Depiction of the late ergosterol pathway in *C. glabrata.* Azoles inhibit lanosterol 14-α-demethylase, the enzyme encoded by *ERG11. NCP1\** and *ERG28\** interact with ergosterol synthesizing enzymes. *ERG29\** is a protein of unknown function involved in ergosterol biosynthesis. Loss of *SET1* results in lower transcript levels of 12 of the 12 ergosterol synthesizing enzymes of the late pathway compared to a wild-type strain upon azole treatment. Genes with decreased transcript levels due the loss of *SET1* are surrounded by a solid square.

**Table S1.**
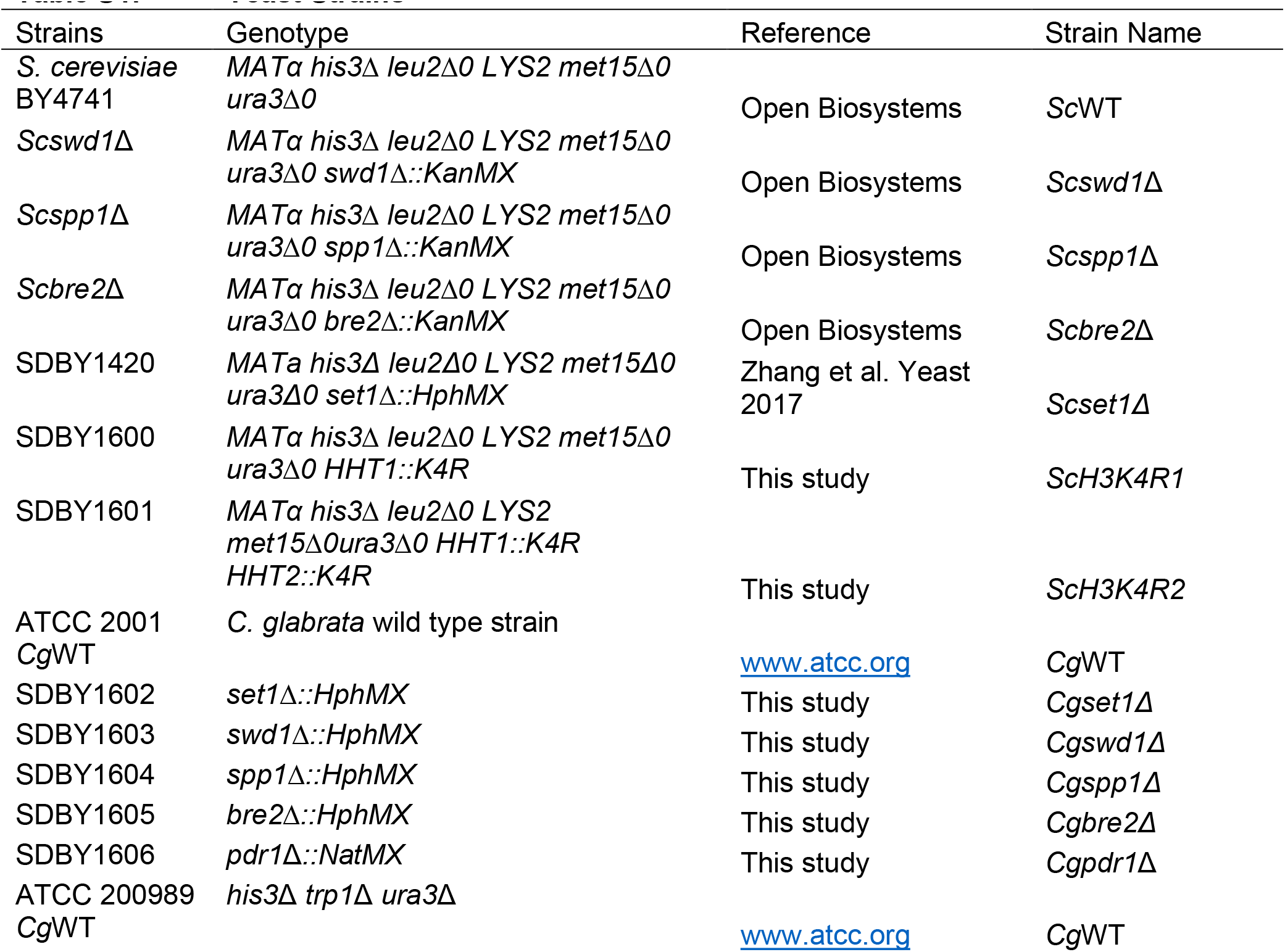
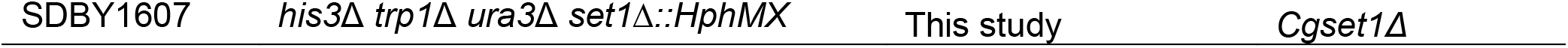
Yeast Strains

**Table S2.** Plasmids

| **Table S2. Plasmids** |  |  |  |  |
| --- | --- | --- | --- | --- |
| **Plasmid** | **Inserted Gene** | **Promoter** | **Vector** | **Source** |
| pGRB2.0 | None |  | pGRB2.0 | Zordan et al. |
| pGRB2.0 | *SET1* | *SET1* | pGRB2.0 | Zordan et al. |
| pGRB2.0 | *SET1/H1048K* | *SET1* | pGRB2.0 | Zordan et al. |

**Table S3.**
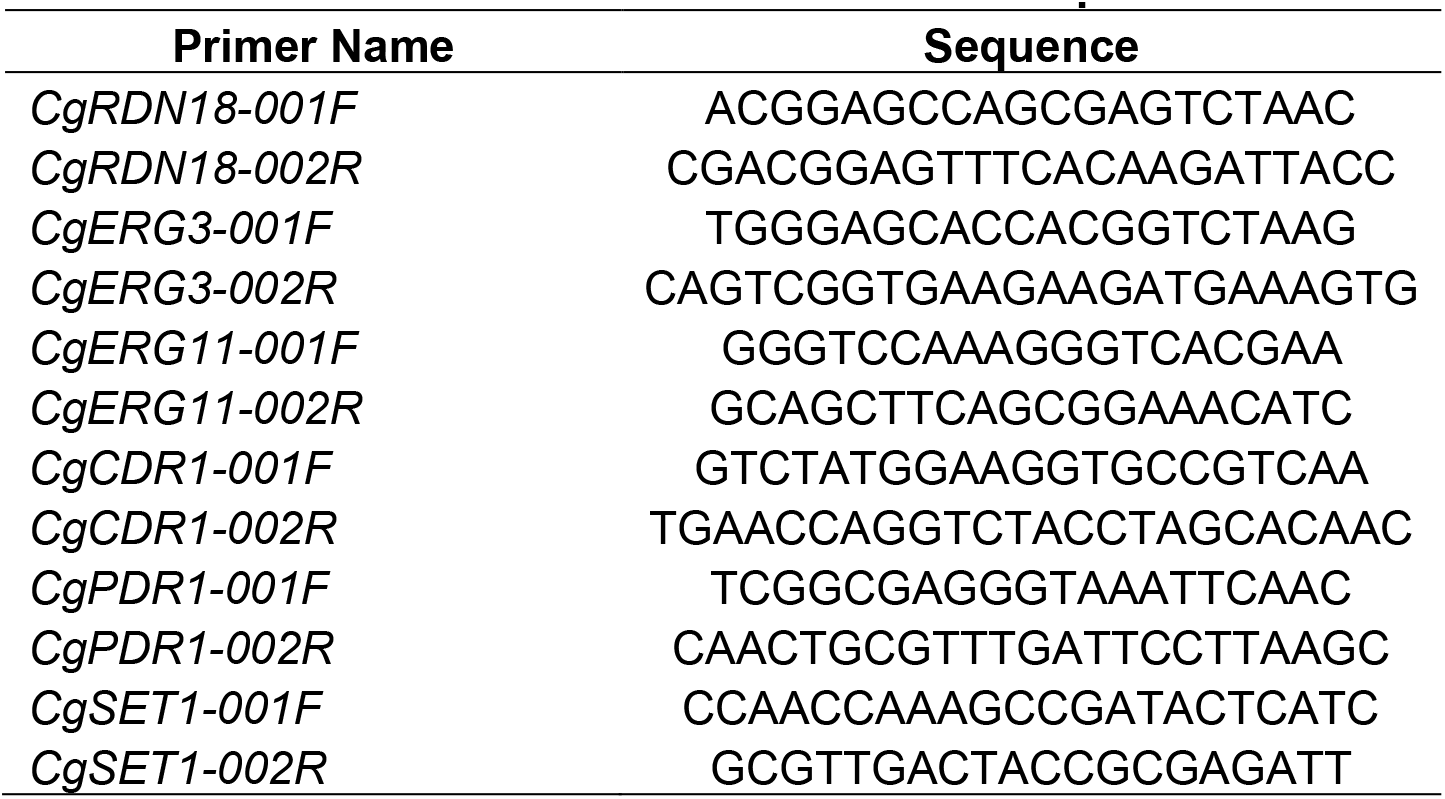
Primers for qRT-PCR

**Table S4.**
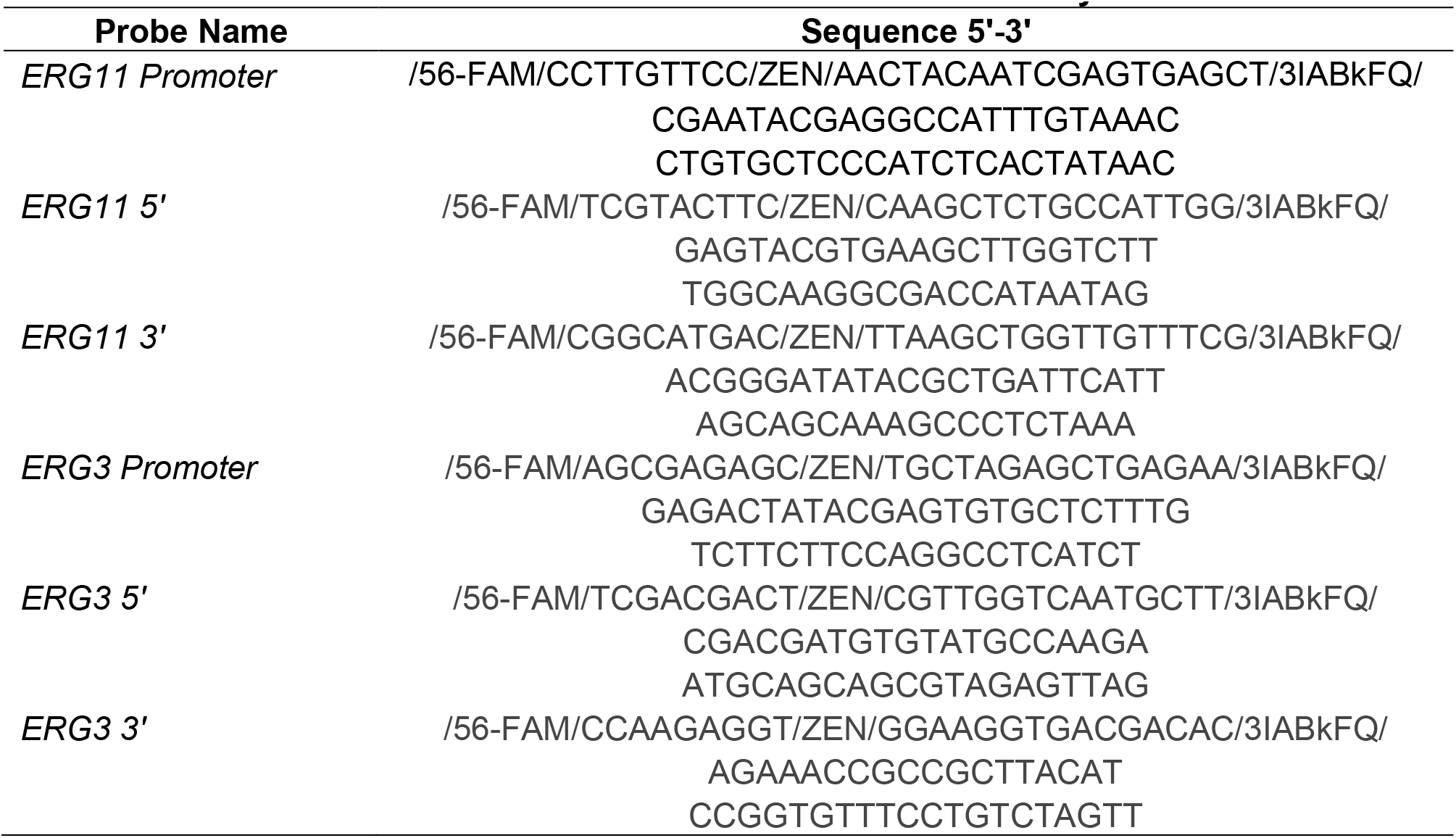
Probe sets for ChIP Analysis

**Table S5.** qRT-PCR Values

| Table S5. |  |  | qRT-PCR Values |  |  |  |
| --- | --- | --- | --- | --- | --- | --- |
| Figure 4A: qRT-PCR |  |  |  |  |  |  |
| Gene | Strain | Condition | Mean RQ | Standard Deviation | N | P-Value |
| *CgCDR1* | *CgWT* | Untreated | 1 | 0.750623 | 7 | n.s. |
| *CgCDR1* | *CgWT* | (+) fluconazole | 4.51 | 1.071572 | 7 | n.s. |
| *CgCDR1* | *Cgset1Δ* | Untreated | 1.1 | 0.972961 | 7 | n.s. |
| *CgCDR1* | *Cgset1Δ* | (+) fluconazole | 4.04 | 0.608125 | 7 | n.s. |
| Figure 4B: qRT-PCR |  |  |  |  |  |  |
| Gene | Strain | Condition | Mean RQ | Standard Deviation |  | P-Value |
| *CgPDR1* | *CgWT* | Untreated | 1 | 0.257724 | 3 | n.s. |
| *CgPDR1* | *CgWT* | (+) fluconazole | 2.29 | 0.204429 | 3 | n.s. |
| *CgPDR1* | *Cgset1Δ* | Untreated | 1.02 | 0.3498374 | 3 | n.s. |
| *CgPDR1* | *Cgset1Δ* | (+) fluconazole | 2.29 | 0.249992 | 3 | n.s. |
| Figure 5E: qRT-PCR |  |  |  |  |  |  |
| Gene | Strain | Condition | Mean RQ | Standard Deviation |  | P-Value |
| *CgERG11* | *CgWT* | Untreated | 1 | 0.32 | 3 | n.s. |
| *CgERG11* | *CgWT* | (+) fluconazole | 10.29 | 0.23 | 3 | n.s. |
| *CgERG11* | *Cgset1Δ* | Untreated | 1.32 | 0.65 | 3 | n.s. |
| *CgERG11* | *Cgset1Δ* | (+) fluconazole | 3.83 | 0.26 | 3 | <0.0001 |
| Figure 5F: qRT-PCR |  |  |  |  |  |  |
| Gene | Strain | Condition | Mean RQ | Standard Deviation |  | P-Value |
| *CgERG3* | *CgWT* | Untreated | 1 | 0.1 | 3 | n.s. |
| *CgERG3* | *CgWT* | (+) fluconazole | 3.63 | 0.69 | 3 | n.s. |
| *CgERG3* | *Cgset1Δ* | Untreated | 1.14 | 0.05 | 3 | n.s. |
| *CgERG3* | *Cgset1Δ* | (+) fluconazole | 0.9 | 0.18 | 3 | <0.01 |
| Figure 6C: qRT-PCR |  |  |  |  |  |  |
| Gene | Strain | Condition | Mean RQ | Standard Deviation |  | P-Value |
| *CgERG11* | *CgWT+V* | Untreated | 0.951 | 0.384 | 4 | n.s. |
| *CgERG11* | *CgWT+V* | (+) fluconazole | 6.791 | 1.988 | 4 | n.s. |
| *CgERG11* | *Cgset1Δ+V* | Untreated | 1.342 | 0.549 | 4 | n.s. |
| *CgERG11* | *Cgset1Δ+V* | (+) fluconazole | 2.603 | 1.043 | 4 | <0.01 |
| *CgERG11* | *Cgset1Δ/SET1* | Untreated | 1.227 | 0.631 | 4 | n.s. |
| *CgERG11* | *Cgset1Δ/SET1* | (+) fluconazole | 6.438 | 2.783 | 4 | n.s. |
| *CgERG11* | *Cgset1Δ/H1048K* | Untreated | 1.346 | 0.429 | 4 | n.s. |
| *CgERG11* | *Cgset1Δ/H1048K* | (+) fluconazole | 3.370 | 0.870 | 4 | <0.05 |
| Figure 6D: qRT-PCR |  |  |  |  |  |  |
| Gene | Strain | Condition | Mean RQ | Standard Deviation |  | P-Value |
| *CgERG3* | *CgWT+V* | Untreated | 1.266 | 1.138 | 4 | n.s. |
| *CgERG3* | *CgWT+V* | (+) fluconazole | 5.686 | 2.144 | 4 | n.s. |
| *CgERG3* | *Cgset1Δ+V* | Untreated | 0.814 | 0.345 | 4 | n.s. |
| *CgERG3* | *Cgset1Δ+V* | (+) fluconazole | 1.512 | 0.429 | 4 | n.s. |
| *CgERG3* | *Cgset1Δ/SET1* | Untreated | 1.034 | 0.64 | 4 | n.s. |
| *CgERG3* | *Cgset1Δ/SET1* | (+) fluconazole | 7.518 | 4.232 | 4 | <0.001 |
| *CgERG3* | *Cgset1Δ/H1048K* | Untreated | 0.888 | 0.206 | 4 | n.s. |
| *CgERG3* | *Cgset1Δ/H1048K* | (+) fluconazole | 2.357 | 0.54 | 4 | <0.05 |

**Figure 6A: ChIP qRT-PCR**
| Gene | Strain | Condition | Mean RQ | Standard Deviation |  | P-Value |
| --- | --- | --- | --- | --- | --- | --- |
| *CgERG11* | *CgWT*<br>promoter | Untreated | 11.24 | 5.568 | 3 | n.s. |
| *CgERG11* | *CgWT*<br>promoter | (+) fluconazole | 22.138 | 12.012 | 3 | n.s. |
| *CgERG11* | *CgWT* 5' | Untreated | 62.531 | 1.452 | 3 | n.s. |
| *CgERG11* | *CgWT* 5' | (+) fluconazole | 133.274 | 20.388 | 3 | <0.001 |
| *CgERG11* | *CgWT* 3' | Untreated | 11.412 | 3.193 | 3 | n.s. |
| *CgERG11* | *CgWT* 3' | (+) fluconazole | 26.019 | 7.85 | 3 | <0.05 |

**Figure 6B: ChIP qRT-PCR**
| Gene | Strain | Condition | Mean RQ | Standard Deviation |  | P-Value |
| --- | --- | --- | --- | --- | --- | --- |
| *CgERG3* | *CgWT* | Untreated | 35.768 | 18.274 | 3 | n.s. |
| *CgERG3* | *CgWT* | (+) fluconazole | 57.996 | 35.319 | 3 | n.s. |
| *CgERG3* | *CgWT* 5' | Untreated | 145.877 | 42.565 | 3 | n.s. |
| *CgERG3* | *CgWT* 5' | (+) fluconazole | 281.392 | 69.117 | 3 | <0.05 |
| *CgERG3* | *CgWT* 3' | Untreated | 50.262 | 13.737 | 3 | n.s. |
| *CgERG3* | *CgWT* 3' | (+) fluconazole | 133.95 | 59.062 | 3 | n.s. |

